# Assessing the Operational Feasibility of Evolutionary Therapy in Metastatic Non-Small Cell Lung Cancer

**DOI:** 10.64898/2026.02.25.707957

**Authors:** Arina Soboleva, Kailas S. Honasoge, Eva Molnárová, Ties A. Mulders, Anne-Marie C. Dingemans, Irene Grossmann, Jafar Rezaei, Kateřina Staňková

## Abstract

Evolutionary cancer therapy (ECT) applies principles of evolutionary game theory to prolong the effectiveness of cancer treatment by curbing the development of treatment resistance. It was shown to increase time to progression while decreasing the cumulative drug dose. ECT individually tailors treatment schedules for patients based on their cancer dynamics and, thus, requires regular follow-up and precise measurements of the cancer burden. The current literature on ECT often overlooks clinical realities, such as rather long intervals between tests, possible appointment delays and measurement errors, in the development of the treatment protocols. In this study, we assess the clinical feasibility of ECT for metastatic non-small cell lung cancer (NSCLC). We create virtual patients with cancer dynamics described by the polymorphic Gompertzian model, based on data from the START-TKI clinical trial. We assess the effects of longer test intervals, measurement error and appointment delays on the expected time to progression under the evolutionary therapy protocols. We show that a higher containment level, although it increases time to progression in the model’s predictions, may lead to premature treatment failure in the presence of measurement error and appointment delay. Further, we show that the ECT protocol with a single containment bound is more robust to the clinical realities than the protocol with two bounds. Finally, we show that a dynamically adjusted treatment protocol can be beneficial for individual patients, but requires a thorough follow-up. This study contributes to the design of a clinical trial and the future clinical implementation of evolutionary therapy for NSCLC.

## 1 Introduction

Evolutionary cancer therapy (ECT), also known as adaptive therapy, applies principles of evolutionary game theory to optimize treatment in metastatic cancer (Gatenby, 2009; Gatenby et al., 2009b; Stanková et al., 2019; West et al., 2020; Wölfl et al., 2021; West et al., 2023). It presents an alternative to the current standard of care in metastatic cancer, which aims at complete eradication of cancer applying maximum tolerable dose (MTD), either continuously or in repeated pre-defined treatment cycles. In metastatic cancer, eradication is rare, and the MTD approach typically promotes rapid development of treatment resistance, causing therapy failure and subsequent progression (Gatenby et al., 2009b; Gatenby, 2009; Dujon et al., 2021; Staňková, 2019; West et al., 2023). ECT typically delays or forestalls the development of treatment-induced resistance by exploiting competition between different types of cancer cells those sensitive and resistant to treatment in the treatment design (Gatenby et al., 2009b; Staňková, 2019; West et al., 2023). ECT is typically guided by evolutionary game-theoretic models of tumor growth calibrated with patient data (Zhang et al., 2022; Strobl et al., 2023; West et al., 2023; Jansén-Storbacka et al., 2026; Beckman et al., 2020). Single-treatment evolutionary therapies typically involve lower doses of treatment than MTD and/or periods without treatment. Because the cumulative dose of ECT is lower than that of MTD, ECT usually has lower toxicity and a higher patient quality of life than standard of care (Zhang et al., 2017, 2022; Cunningham et al., 2018).

The first ECT clinical trial, a small Zhang et al.’s trial in metastatic castrate-resistant prostate cancer (mCRPC), was initiated in 2015 (Zhang et al., 2017, 2022). The patients were treated with MTD of the drug Abiraterone until the tumor burden measured through prostate-specific antigen (PSA), a biomarker used as a proxy for overall tumor burden, decreased to 50% of its initial level. At that point, the treatment was paused to allow the tumor burden to regrow. When the initial level of PSA was reached again, the treatment was resumed and a new treatment cycle began (Zhang et al., 2017). As a result of the clinical trial, time to progression (TTP) increased to 33.5 months compared to 14.3 months in the standard of care group, while the cumulative drug dose was decreased by half (Zhang et al., 2022). The success of this initial trial led to a growing interest in evolutionary cancer therapy with multiple studies published on theoretical analysis of (potential) mathematical models that could guide ECT (Viossat & Noble, 2021; Alvarez & Viossat, 2024), validation of these models with real-world data (Kaznatcheev et al., 2019; Ghaffari Laleh et al., 2022; Soboleva et al., 2025b; Garjani et al., 2025), development of different ECT approaches, including dose stabilization (Hockings et al., 2023; Salvioli et al., 2024; Cunningham et al., 2020b), intermittent dosing (Yu et al., 2017; De Iuliis et al., 2016), extinction therapy (Gatenby et al., 2019), and the double-bind therapy (Gatenby et al., 2009a; Basanta et al., 2012). Moreover, evolutionary cancer therapy is being evaluated in an increasing number of clinical trials across cancers, including prostate (NCT05393791), ovarian (NCT05080556), melanoma (NCT03543969), and skin cancers such as basal cell carcinoma (NCT05651828).

Many studies have applied optimal control and related optimization techniques to evolutionary game theoretic (EGT) models, as well as to other mathematical models of eco-evolutionary cancer dynamics under treatment, to derive therapeutic protocols that delay resistance, prolong time to progression, and improve patient quality of life (Martin et al., 1992; Tomlinson, 1997; Carrère, 2017; Cunningham et al., 2018; Brady-Nicholls & Enderling, 2022; Wang & Lei, 2025; Gluzman et al., 2020; Stanková et al., 2019) These efforts are part of the broader and rapidly expanding field of mathematical oncology, in which optimization and evolutionary modeling are increasingly integrated into clinical decision-making (Scibilia et al., 2025; Pradelli et al., 2026; Soboleva et al., 2025a). Such approaches span a spectrum from purely theoretical analyses to optimization of models calibrated and validated against experimental and/or clinical data.

An alternative to Zhang et al.’s protocol is the containment strategy. Unlike the original approach, which maintains tumor burden within two predefined bounds, the containment protocol relies on a single threshold. Treatment is suspended once the tumor burden decreases below this threshold and resumed when it exceeds it again. The key challenge is, therefore, how this containment level should be optimally determined. It has been theoretically shown that a higher containment level can prolong time to progression (Viossat & Noble, 2021). However, this potential benefit comes with risk, as a higher tumor burden may increase the probability of new mutations, the likelihood of metastatic spread, and the severity of patient symptoms, thereby reducing quality of life (Joseph et al., 2018; He et al., 2025; Lommen et al., 2026; Howard et al., 2011). To address this trade-off, Gallagher et al. derived the containment threshold using competition-based tumor growth models, requiring that under discrete monitoring intervals the tumor cannot grow from the containment level to the progression threshold between successive tumor measurements (Gallagher et al., 2025).

The applications of ECT for metastatic non-small cell lung cancer (NSCLC) are currently being evaluated (Jansén-Storbacka et al., 2026), based on fitting *in vitro* and *in vivo* data to mathematical models (Kaznatcheev et al., 2019; Ghaffari Laleh et al., 2022; Soboleva et al., 2025b; Garjani et al., 2025). The state-of-the-art treatment for metastatic NSCLC patients with known driver mutations is targeted therapy with tyrosine kinase inhibitors (TKIs). In epidermal growth factor receptor (*EGFR*)-mutated NSCLC, third-generation TKIs such as osimertinib significantly improve progression-free and overall survival compared to earlier-generation inhibitors (Ramalingam et al., 2020). However, despite these advances, resistance inevitably emerges and tumor regrowth occurs. Clinical and molecular analyses of patients progressing on *EGFR* TKIs reveal substantial heterogeneity in resistance mechanisms, including secondary *EGFR* alterations, MET amplification, and alternative bypass pathways, reflecting the dynamic evolutionary nature of the disease (van der Wel et al., 2025; Steendam et al., 2020; Rolfo et al., 2021; Chmielecki, 2023).

Prospective studies combining tumor tissue and circulating tumor DNA analyses further demonstrate that single-modality testing may miss clinically relevant resistance mechanisms, underscoring spatial and temporal heterogeneity during treatment (van der Wel et al., 2025; Ernst et al., 2025). These findings support the view that resistance development in metastatic NSCLC is an evolutionary process occurring within a heterogeneous tumor ecosystem (Gatenby & Vincent, 2003), a perspective that has motivated both eco-evolutionary modeling and optimization-based approaches to adaptive therapy.

From a theoretical perspective, mathematical models of tumor growth and resistance evolution suggest that continuous maximum tolerated dose therapy may promote competitive release of resistant cell populations. In contrast, containment-based strategies aim to maintain a controlled tumor burden by preserving a population of treatment-sensitive cells that suppress the expansion of resistant phenotypes. Analytical and computational studies of competition-based tumor growth models demonstrate that containment protocols can prolong time to progression across a broad range of biologically plausible parameter regimes (Viossat & Noble, 2021; Alvarez & Viossat, 2024). Recent data-driven modeling of longitudinal NSCLC tumor burden, incorporating density-dependent growth and frequency-dependent interactions, further indicates that evolutionary therapy protocols may theoretically extend progression-free survival compared to continuous treatment (Jansén-Storbacka et al., 2026).

However, clinical implementation of ECT for metastatic NSCLC is challenging because it requires careful monitoring of disease progression (Soboleva et al., 2025a). Unlike prostate cancer, NSCLC does not have an easily measured serum biomarker, and the disease is usually monitored through imaging using computed tomography (CT), brain magnetic resonance imaging (MRI) and total body positron emission tomography (PET) scans (Molnárová et al., 2025). The lesions on the images are often difficult to measure, which leads to high variation of the estimated tumor volume and potential disagreements between measurements made by different radiologists (Oxnard et al., 2011b; Zhao et al., 2009).

Another challenge is the long interval between assessments. In current clinical practice, radiological response evaluation in metastatic NSCLC is typically performed every 6–12 weeks (Hendriks et al., 2023). In evolutionary therapy, where patients may have treatment vacations, such long intervals can be risky because clinically relevant regrowth may occur between scans. Moreover, in routine care, imaging and follow-up appointments may be delayed (e.g., due to scheduling constraints or patient-related factors), further increasing the effective interval between measurements (Iwsakul & Geater, 2025).

In this study, we evaluate the feasibility of the common evolutionary therapy protocols Zhang et al.’s, containment and dynamic containment protocols - in clinically realistic conditions through simulations. Based on time-series volumetric patient data from the START-TKI trial (NCT05221372), we utilize frequency-dependent polymorphic Gompertzian model to simulate virtual patients corresponding to the patient cohort of metastatic NSCLC treated with tyrosine kinase inhibitor osimertinib. For these virtual patients, we compare time to progression under the considered evolutionary therapy protocols and the continuous MTD treatment under assumptions of varying measurement errors and appointment delays. We then summarize our findings as suggestions for the best evolutionary osimertinib protocol for metastatic NSCLC, accounting for the clinical realities.

## 2 Methods

### 2.1 The frequency-dependent Gompertzian model

We assumed that cancer eco-evolutionary dynamics follow the polymorphic frequency-dependent Gompertzian model with competition between sensitive and resistant cancer cell populations and log-kill osimertinib-induced death rate. The model was selected as it demonstrated the highest accuracy when fitted to the NSCLC patient data in previous studies (Soboleva et al., 2025b; Jansén-Storbacka et al., 2026). The model describes the eco-evolutionary dynamics of treatment-sensitive and treatment-resistant cancer cell populations denoted by *S*(*t*) and *R*(*t*), respectively, as follows:

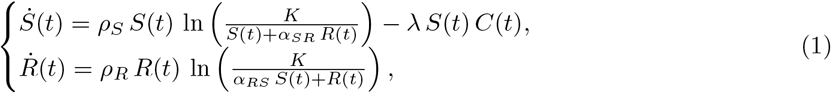

where *ρ*_*S*_ and *ρ*_*R*_ are growth rates of sensitive and resistant populations, respectively, *K* is carrying capacity representing cancer cell population limit to growth due to space and resource constraints, *α*_*RS*_ and *α*_*SR*_ capture the competitive effect of sensitive population on the resistant one and vice versa, respectively, *λ* reflects how strongly treatment affects sensitive population, and *C*(*t*) indicates the drug dose at time *t*. The total cancer population *N* (*t*) at time *t* is defined as *N* (*t*) = *S*(*t*) + *R*(*t*).

### 2.2 Virtual patients

The frequency-dependent Gompertzian model was parametrized using patient data from 37 patients with metastatic NSCLC treated with first-line tyrosine kinase inhibitor osimertinib from the STARTTKI trial (NCT05221372). The dataset comprises longitudinal volumetric measurements extracted from three-dimensional imaging, with CT scans used for lung lesions and MRI for brain metastases. The tumor burden *N* (*t*) at every measured point was estimated as the sum of volumes of all measured lesions. We fitted parameters *ρ*_*S*_, *ρ*_*R*_, *α*_*SR*_, *α*_*RS*_, *K*, and *λ* of the frequency-dependent polymorphic Gompertzian model (1) individually to each patient’s tumor burden trajectory by minimizing the mean squared error (MSE), obtaining one patient-specific parameter set per patient (37 in total). The empirical distribution of the fitted parameters was used to estimate a multivariate normal distribution, from which virtual patient parameter sets were sampled. These simulated patients were then used to compare containment-based evolutionary strategies with MTD therapy in terms of time to progression, defined as the first occurrence of total tumor burden exceeding 1.2*N* (0) (a 20% increase over baseline). In this framework, treatment effects were modeled exclusively through the treatment variables *C*(*t*) and *λ*, while intrinsic growth and interaction parameters were assumed to be independent of the dosing schedule. This assumption allows us to simulate alternative treatment approaches and compare them to MTD. More information on the data and the fitting procedure is available at Appendix A1.

Based on the 37 parameter sets, we constructed a multivariate normal distribution from which we sampled 100 virtual patients, each representing a cancer dynamics described by an individually parametrized polymorphic Gompertzian model. We simulated the treatment dynamics by solving the ODE system 1 with the Runge–Kutte 45 method from the `solve_ivp` module in the `scipy.integrate` library in Python. To ensure realism of the virtual patients’ dynamics, we only included the once with predicted time to progression under MTD protocol (*C*(*t*) = 1 ∀ *t*) below 3000 days. Furthermore, we did not include non-responders - virtual patients whose tumor size had not decreased below half of the initial size in the first 120 days of treatment. Using virtual patients allowed us to explore a wider range of potential tumor dynamics.

### 2.3 Tested treatment protocols

In our study, we tested different evolutionary therapy protocols and assessed the outcomes in terms of time to progression. We defined progression as an increase in tumor burden by more than 20% of its initial tumor burden, following previous modeling studies (Viossat & Noble, 2021; Strobl et al., 2021; Gallaher et al., 2018). We considered binary on–off treatment protocols, in which the dose is fixed at MTD when administered and zero otherwise. In the model (1), the medication variable is *C*(*t*) ∈ {0, 1}, where *C*(*t*) = 1 corresponds to the standard osimertinib dose of 80 mg. In the treatment protocols considered, the decision to stop or resume medication can be changed daily, reflecting the clinical reality of osimertinib administered as a daily pill. We assume that the test that measures tumor burden and the doctor appointment, at which the decision to stop or resume treatment is made, occur on the same day. With longer test intervals there is a possibility of premature failure of the treatment protocol. Progression is considered premature if the tumor burden exceeds the progression threshold during the treatment holidays. In this case, the sensitive cancer population may still be present and the medication can still be effective; however, progression is declared, according to the predefined 20% increase from the initial tumor burden.

Let 𝒯= {*t*_0_, *t*_1_, …} denote the discrete testing times with *t*_0_ = 0 and *t*_*i*+1_ *> t*_*i*_. Define the (continuous) time of progression as *τ*_prog_ = min{*t* ≥ 0 : *N* (*t*) *> N*_prog_}. Progression is observed at *t*_*K*_ = min{*t*_*i*_ ∈ 𝒯 : *t*_*i*_ ≥ *τ*_prog_}. In simulations, treatment decisions are updated only at testing times and the simulation is terminated once progression is observed at *t*_*K*_. We report *τ*_prog_ as the time to progression to avoid the discretization artifact of the longer test intervals. In the simulations, we consider the following treatment protocols:

- **Continuous MTD treatment**. The current standard of care. Under this protocol, patients receive the constant treatment dose daily until progression. In the model, it is reflected as

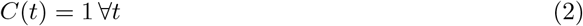
- **Zhang et al.’s evolutionary protocol**. Patients receive a constant treatment dose until the tumor burden reaches the established lower threshold *N*_min_ (in the initial clinical trials *N*_min_ = 0.5 *N* (0) (Zhang et al., 2017, 2022)). The treatment is then paused until the tumor burden regrows to the upper threshold *N*_max_, at which point it is resumed. This is repeated until progression. At each testing time point *t*_*i*_, the treatment is set for the next interval between tests (*t*_*i*_, *t*_*i*+1_] according to

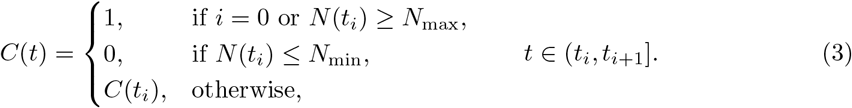
- **Containment protocol**. Unlike in Zhang et al.’s protocol, this protocol has a single threshold *N*_cont_. The patient is treated at a constant dose until the tumor burden decreases below *N*_cont_; then, treatment is paused. The treatment is resumed when the tumor size exceeds *N*_cont_. At each testing point *t*_*i*_, the treatment for the next interval is set as

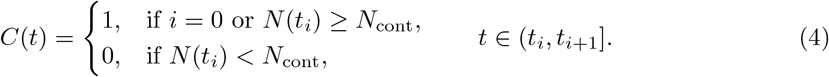

### 2.4 Effect of appointment delay and measurement error

We incorporated appointment delays into the simulations to reflect clinical reality, where appointments are often postponed due to the doctor’s scheduling conflicts, vacations, public holidays, worsening of the patient’s condition, etc. The time to the next test was determined as the interval specified in the treatment protocol plus an additional delay sampled from a Poisson distribution (in days). The next testing moment is defined as the following:

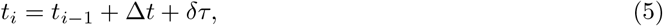

where Δ*t* is the intended interval between tests and *δτ* is a random appointment delay in days, with *δτ* ~ Poisson(*λ*).

The delay was sampled independently at each testing point and was constrained to be non-negative, reflecting that appointments may be postponed but not advanced. We considered different mean values *λ* of the Poisson distribution (*λ* ∈ {2, 5, 7, 10, 15}), up to 15 days, corresponding to half of a standard 30-day testing interval. For each value of *λ*, we ran 1000 simulations to account for the stochastic nature of the delay process and analyzed the resulting distributions of TTP.

To account for measurement uncertainty in radiological assessments, we modeled the observed tumor volume at each testing point as the model-predicted tumor size multiplied by a stochastic error term drawn from a log-normal distribution:

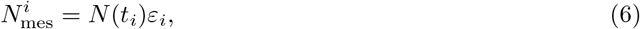

where 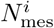 denotes the observed tumor burden at time *t*_*i*_, *N* (*t*_*i*_) is the model-predicted tumor size at the time point *t*_*i*_ and *ε*_*i*_ ~ LogNormal (0, *σ*^2^) represents multiplicative measurement error, sampled from log-normal distribution.

We modeled measurement error as a multiplicative uncertainty relative to the model-predicted tumor size. In the log-normal distribution, we set *µ* = 0 and considered *σ* ∈ {0.02, 0.05, 0.1, 0.2, 0.3}. These values correspond to increasing levels of variability, ranging from a minimal error of 2% to a high variability of 30%. The measurement error can both increase and decrease the actual tumor burden.

Because the error is sampled independently at each measurement time point, its cumulative effect on the treatment trajectory and the resulting TTP is stochastic and varies across simulation runs. Therefore, for each patient and each value of *σ*, we performed 1000 simulations to characterize the distribution of outcomes under measurement uncertainty.

We then assessed the joint effect of measurement error and appointment delay. We assumed error and delay to be independent, and for all combinations of *λ* for the appointment delay and *σ* for the measurement error, we ran 1000 simulations per patient and analyzed the distributions of TTPs.

### 2.5 Dynamically adjusted treatment protocol

We also simulated a dynamic containment protocol in which the containment threshold *N* ^∗^ is updated during treatment based on the tumor burden growth rate observed during medication-free intervals. The objective of this protocol is to maintain the containment level as high as possible to maximize the potential TTP benefit, while ensuring that the tumor burden cannot reach the progression threshold between successive evaluations. The protocol is inspired by the method proposed by Gallagher et al. (Gallagher et al., 2025), who derived the optimal threshold as the maximum level from which the untreated tumor cannot grow to the progression threshold within one testing interval.

In clinical practice, it is not feasible to reliably identify and parametrize a specific tumor growth model from sparse longitudinal measurements. Therefore, we employ a linear growth estimate to avoid model-specific assumptions and to assess the feasibility of dynamic containment under realistic clinical conditions.

To reduce the risk of premature failure due to growth-rate misestimation, we restrict the containment threshold to *N* ^∗^ ∈ [0.4 *N* (0), 0.8 *N* (0)]. The containment threshold cannot exceed 80% of the initial tumor burden to avoid an erroneous growth rate estimate, leading to a too high containment threshold and premature failure. It also cannot be lower than 40% of the tumor burden, as it increases the risk of resistance development and would undermine the concept of evolutionary therapy. The containment threshold *N* ^∗^ is determined at the end of each treatment-free interval during which the tumor burden increased as

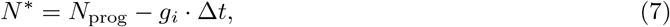

where Δ*t* is the established interval between tests (not accounting for the appointment delay) and *g*_*i*_ is the linear growth rate estimated at the *i*th measurement time point, calculated as

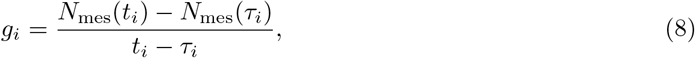

where *t*_*i*_ is the time of the *i*th measurement, *N*_mes_(*t*_*i*_) is the corresponding measured tumor burden, *τ*_*i*_ is the time of the last treatment pausing decision prior to *t*_*i*_, *τ*_*i*_ = max {*t*_*j*_ : *t*_*j*_ *< t*_*i*_ and *C*(*t*_*j*_) = 1)} and *N*_mes_(*τ*_*i*_) is the tumor burden measurement at this point. Note that we used the measured tumor burden *N*_mes_(*t*_*i*_), which may be subject to measurement error, rather than the model-predicted value *N* (*t*_*i*_).

As with the previously introduced protocols, we incorporated appointment delays and measurement error into the dynamic containment protocol and assessed their impact on TTP. The virtual patients’ data, simulations, and the analysis code implemented in Python 3.12.3 are available at: https://gitlab.tudelft.nl/evolutionary-game-theory-lab/clinical-feasibility.

## 3 Results

### 3.1 Trade-off between protocol threshold levels and test interval length

We first simulated an idealized scenario of daily testing. Figure 1A illustrates the tumor burden dynamics under MTD, Zhang et al.’s protocol, and the containment protocols in such settings. The on/off medication cycles are repeated until progression, defined as the first time when the tumor burden exceeds 1.2 *N* (0). Under Zhang et al.’s protocol, the patient undergoes extended periods of treatment and treatment holidays, whereas under the containment protocol, treatment is adjusted almost daily. Both Zhang et al.’s protocol and the containment protocol result in a substantial increase in TTP compared to continuous MTD treatment. This is consistent with the theoretical results of Viossat and Noble, who showed that both strict containment and intermittent containment (containment between two bounds, as in Zhang et al.’s protocol) outperform MTD in terms of time to progression for cancer dynamics described by the Gompertzian model (Viossat & Noble, 2021).

**Figure 1.**
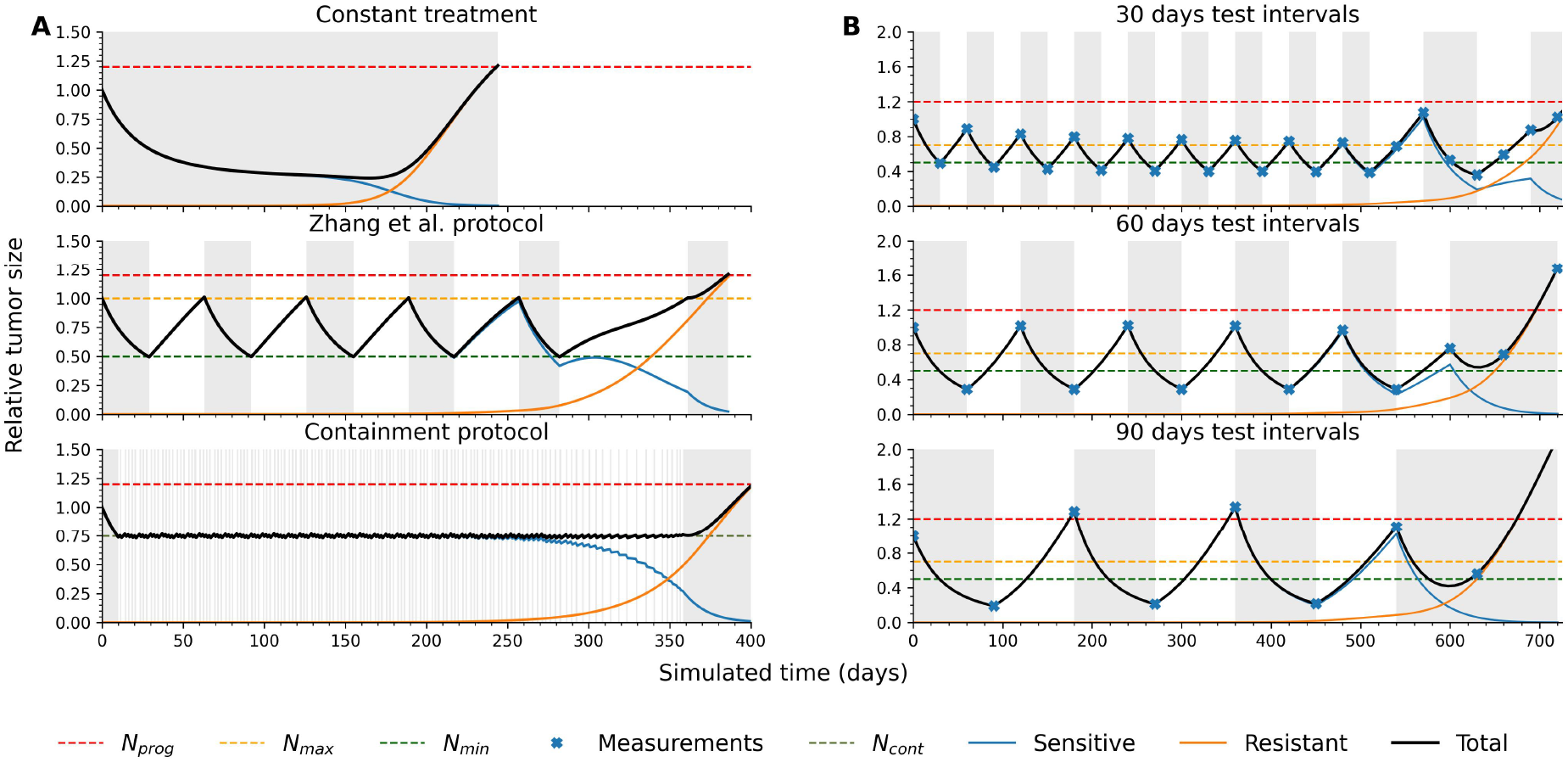
The cancer dynamics under constant treatment, Zhang et al.’s protocol and containment protocol with 30−, 60− and 90− day intervals between tests. Blue, orange and black lines indicate the dynamics of sensitive, resistant, and total cell populations, respectively. The yellow and green dotted lines show *N*_max_ and *N*_min_ levels of Zhang et al.’s protocol. The red dashed line indicates the progression threshold, denoted 1.2 *N* (0). When the tumor burden reaches this threshold, the treatment protocol has failed. The gray areas indicate the periods during which the treatment was in effect. **A.** For all three protocols in these simulations, the decisions are made daily. Both Zhang et al.’s and containment protocols significantly increase time to progression compared with the constant-treatment protocol. The containment protocol yields a slightly higher TTP than Zhang et al.’s protocol, due to the higher containment level. **B**. Blue crosses indicate the points of measurement. The 90-day test interval eventually leads to premature treatment failure.

We varied the upper bound *N*_max_ in Zhang et al.’s protocol and the containment level *N*_cont_ in the containment protocol. The TTP benefit was observed across all patients and threshold levels, and was more pronounced at higher thresholds. For conservative bounds of *N*_max_ = 0.7 in Zhang et al.’s protocol and *N*_cont_ = 0.5 in the containment protocol, the TTP benefit across patients ranged from 1.02 to 37.6 (median 2.07) under Zhang et al.’s protocol and from 1.01 to 37.6 (median 1.69) under the containment protocol. The increase in TTP varied substantially between patients; however, even when it was modest, patients could still benefit from reduced medication exposure and the associated reduction in side effects. The patients were off medication for, on average, 40% of the treatment trajectory under Zhang et al.’s protocol and 33% under the containment protocol.

As daily follow-up is unrealistic in clinical practice, we simulated treatment protocols under monthly (30 days), bimonthly (60 days), and trimonthly (90 days) tumor assessments (Figure 1B). In such settings, the tumor volume can decrease significantly below the lower bound of Zhang et al.’s protocol during the treatment period, and significantly exceed the upper bound during the treatment holidays. Under the containment protocol, the cycling of the tumor burden also becomes less frequent, making it qualitatively similar to Zhang et al.’s protocol under clinically realistic monitoring intervals.

The main risk of longer test intervals is the premature failure of the treatment protocol, defined as the tumor burden exceeding the progression threshold during the treatment holidays (Figure 1B with the 90-day test interval). The percentage of premature failures across virtual patients depends both on the intervals between tests and bounds of evolutionary therapy protocols. However, even with low containment bound of *N*_cont_ = 0.5 *N* (0), a 60-day test interval leads to premature failure for 78% of patients, and a 90-day interval in 90% of patients. We therefore fixed the interval between tests to 30 days and varied the bounds of the evolutionary therapy protocols. The increase in TTP at higher protocol thresholds is modest, whereas the number of premature failures increases substantially (Appendix A2). We therefore continue the analysis using Zhang et al.’s protocol with *N*_min_ = 0.5 *N* (0) and *N*_max_ = 0.7 *N* (0), and the containment protocol with *N*_cont_ = 0.5 *N* (0).

With the established test interval and protocols’ bounds, 44 out of 100 virtual patients demonstrated premature failure under both Zhang et al.’s and containment protocols. Essentially, those cases demonstrated growth from 0.5*N* (0) to 1.2*N* (0) in less than 30 days. In real clinical practice, such growth would be very unlikely. We therefore excluded these 44 cases from further analysis. Additionally, three patients experienced premature failure with Zhang et al.’s protocol but not with the containment protocol. This can be explained by Zhang et al.’s protocol’s upper bound being higher than the containment threshold, making it riskier with longer test intervals. Furthermore, 15 virtual patients reached a TTP of 5000 days, corresponding to the simulation time limit. These patients exhibited unrealistically slow resistance development and were therefore excluded from further analysis. The parameter sets leading to unrealistic dynamics arise from limitations in the clinical data and fitting procedure. Because the virtual patients were sampled from the distribution inferred from the fitted patient data, they inherit these artifacts (more details in Appendix A1). The subsequent analyses were performed using the remaining 38 virtual patients. These patients exhibit disease dynamics consistent with clinical observations and provide additional variation in regrowth rates, regrowth patterns, and TTP under continuous treatment. We therefore continued the analysis with this cohort. This also maintains a manageable sample size for analysis and visualization.

For 27 of the 38 virtual patients, the TTP under Zhang et al.’s protocol exceeds that under the containment protocol with a 30-day test interval. This difference is attributable to the higher tumor burden permitted in Zhang et al.’s protocol (*N*_max_ = 0.7 *N* (0)), which is associated with longer TTP (Viossat & Noble, 2021).

### 3.2 Effect of appointment delay

Appointment delays can alter the treatment trajectory of evolutionary therapy protocols and lead to a decrease in the TTP. Figure 2A-B shows the principles of how a delayed testing moment can reduce TTP. Delays during the off-medication period are risky for patients with rapidly progressing cancer, as they may lead to premature failure within the same treatment cycle (Figure 2A). A delay during the on-medication period does not cause immediate protocol failure; rather, the prolonged on-medication interval eliminates more sensitive cells, thereby allowing the resistant population to proliferate (Figure 2B). As a result, the tumor becomes resistant earlier, and TTP is shorter. In specific cases, appointment delays may be beneficial by allowing the tumor burden to rise above or fall below the protocol bounds. In this way, the delay can prevent an additional treatment holiday that would otherwise lead to premature failure, or an additional treatment interval that would accelerate the development of resistance (Appendix A3).

**Figure 2.**
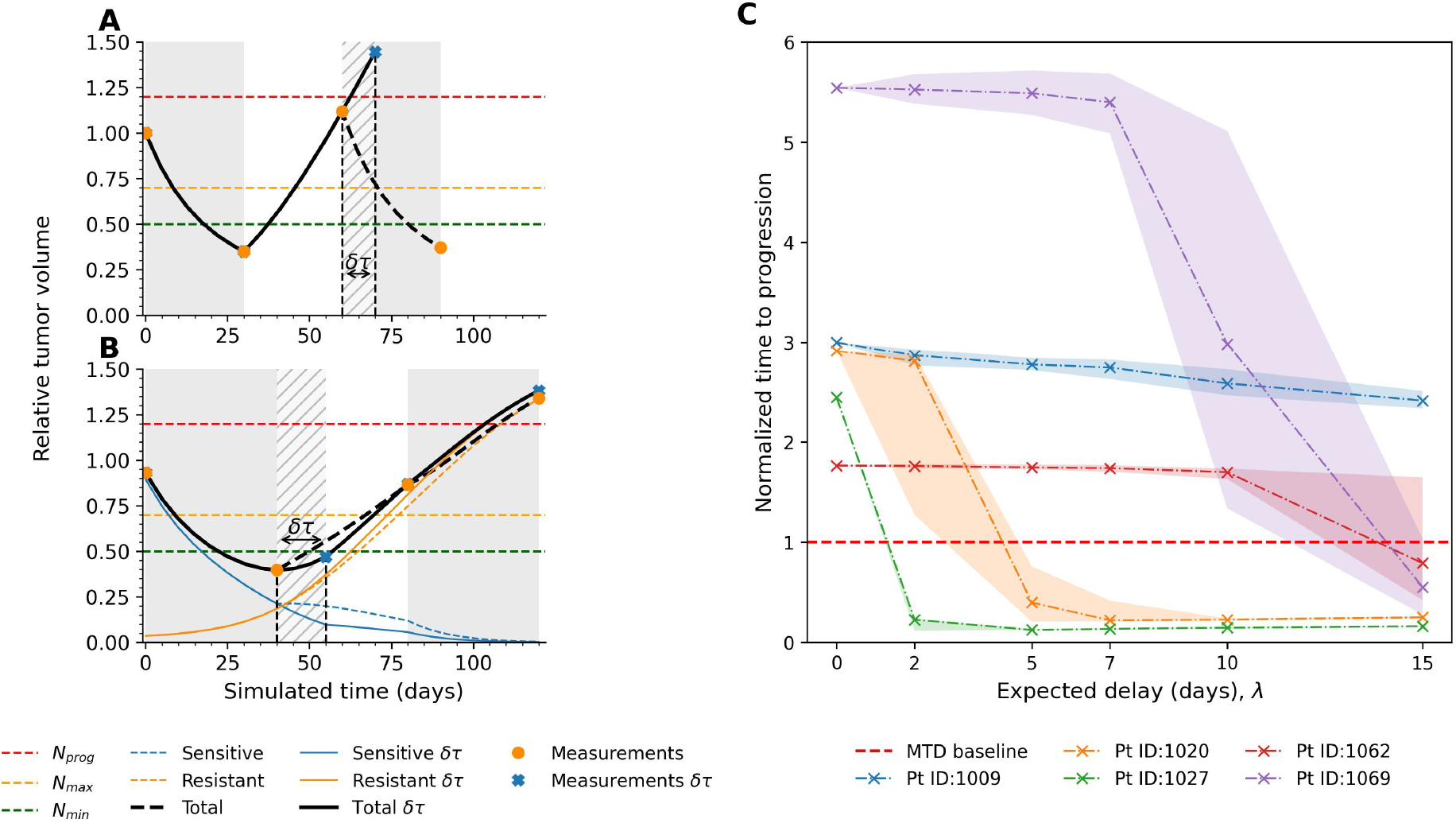
Effect of appointment delay on TTP under Zhang et al.’s protocol. **A-B** Red, yellow, and green dashed horizontal lines demonstrate the progression threshold, upper and lower bounds of Zhang et al.’s protocol, respectively. The black line shows the total tumor burden up to the occurrence of the delay and, thereafter, the tumor dynamics under delayed assessment; the dashed line shows the corresponding dynamics without delay. The orange circles indicate the measurement points along the treatment trajectory without delay, whereas the blue crosses indicate those with delay. The dashed blue and orange lines show the dynamics of the sensitive and resistant populations, respectively, under the treatment trajectory without delay, whereas the solid lines show the same under the trajectory with delay. **A** The delay of appointment during the off-medication period leads to a premature failure of the treatment protocol as the resuming of treatment is postponed. **B** The delay during the on-medication period leads to “overtreatment”, resulting in excessive eradication of sensitive cells and consequently earlier emergence of resistance and progression. **C** Medians of TTPs of 1000 simulations across increasing expected delay *λ*, normalized to the TTP under continuous treatment for the same patient. The crosses show the median TTPs for each value of *λ* and the shaded areas show the interquartile range. The tumor dynamics across virtual patients exhibit varying sensitivity to appointment delay. For example, patient 1009 shows only a slight decrease in median TTP, never falling below the MTD baseline. Patients 1062 and 1069 are robust to small expected delays, but TTP decreases substantially for delays exceeding one week. In contrast, patients 1020 and 1027 are highly sensitive, as even small expected delays lead to premature protocol failure.

Due to the stochastic nature of the delay in the simulations, TTP across simulation runs can vary significantly. We report the median TTP and interquartile range (IQR) for each patient and each *λ* value to summarize central tendency and variability. The median and IQR were chosen over the mean and standard deviation because the distributions of TTPs are often non-normal and skewed. For the majority of virtual patients (24 of 38 under Zhang et al.’s protocol and 21 of 38 under the containment protocol), the median TTP decreased with increasing expected delay *λ*. For the remaining virtual patients, the median TTP either fluctuated or remained stable across values of the expected delay *λ*. The magnitude of the decrease in median TTP differs substantially across patients. Figure 2C illustrates the decrease in median TTP under Zhang et al.’s protocol as the expected delay *λ* increases for five representative patients. Patients 1027 and 1020 are highly sensitive to delays, as even an expected delay of 2 or 5 days leads to premature failure of Zhang et al.’s protocol. For patients 1069 and 1062, Zhang et al.’s protocol is robust to short expected delays of up to a week; however, the median TTP falls below the MTD baseline for larger values of *λ*. Finally, for patient 1009, the median TTP decreases only slightly and does not fall below the MTD baseline. This suggests that the risk of premature failure is low, and that the modest reduction in TTPs is attributable to earlier emergence of resistance caused by prolonged periods of treatment exposure. No substantial difference was observed in the sensitivity of Zhang et al.’s and the containment protocols to appointment delays. In the majority of cases, the median TTP under Zhang et al.’s protocol remains higher than that of the containment protocol across all tested values of *λ* (Figure A3).

### 3.3 Effect of measurement error

Measurement error can affect treatment decisions at the testing time points by causing the measured tumor burden to fall above or below the protocol bounds. Figure 3 describes the ways measurement error can affect the treatment trajectory. For example, if the actual tumor burden is slightly above *N*_max_ (or *N*_cont_ for the containment protocol), so that treatment should be resumed, measurement error may yield an observed value below the threshold. This can result in an additional treatment holiday and potentially premature failure (Figure 3B). If the true cancer burden is close to the progression threshold, an error making the measurement larger can lead to a premature declaration of treatment failure (Figure 3A). Further, an erroneous measurement can be higher than *N*_min_ (or *N*_cont_), leading to an additional interval on medication, even though the actual tumor burden is below the threshold (Figure 3C). Unnecessary treatment periods in these cases promote the development of resistance.

**Figure 3.**
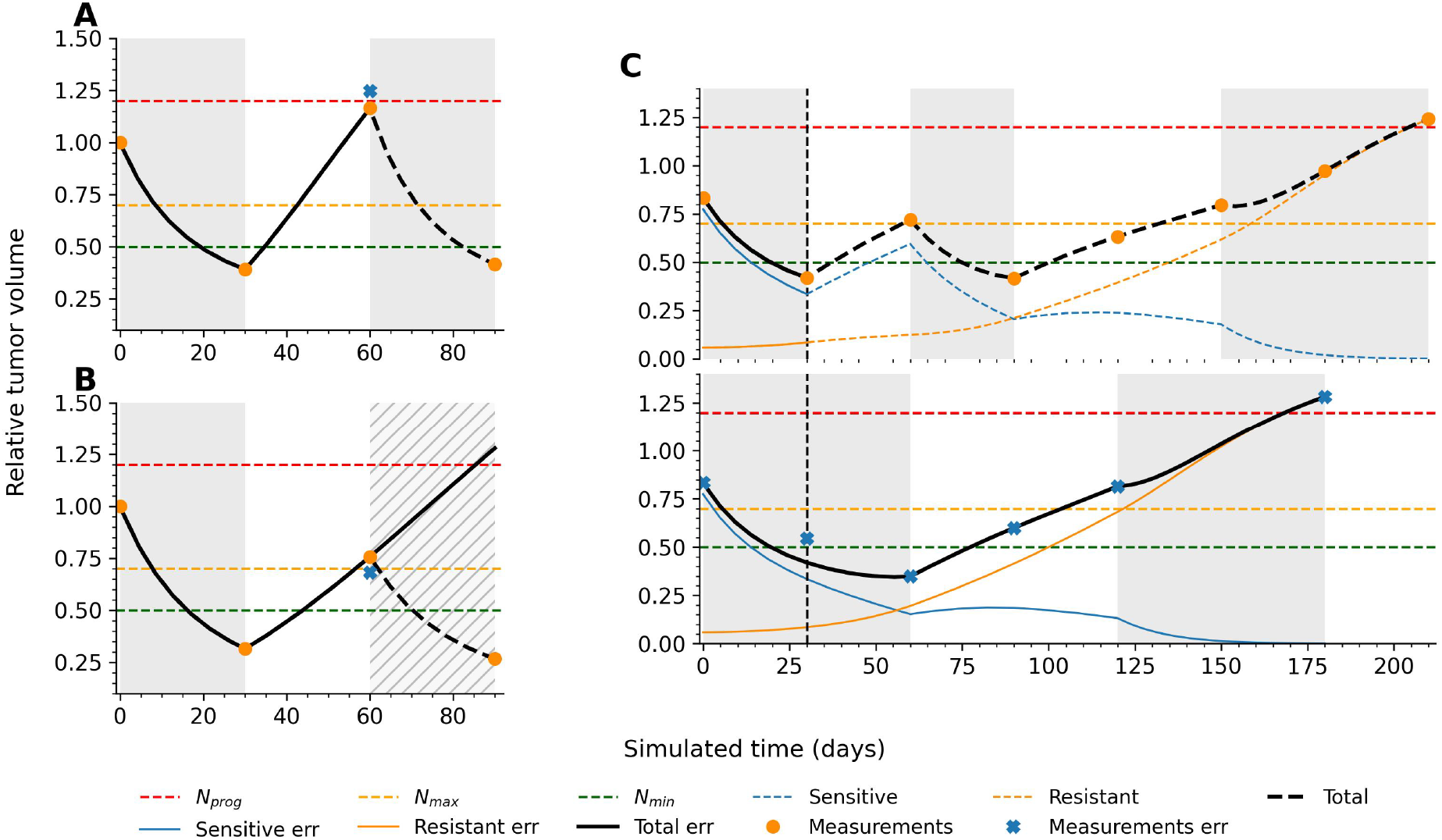
The logic of measurement error affecting TTP of Zhang et al.’s protocol. Red, yellow, and green dashed horizontal lines demonstrate the progression threshold and upper and lower bounds of Zhang et al.’s protocol, respectively. The black line shows the total tumor burden up to the occurrence of the measurement error and, thereafter, the dynamics with error; the dashed line shows the corresponding dynamics without error. The orange circles indicate measurement points along the treatment trajectory without error, whereas the blue crosses indicate those with error. The dashed blue and orange lines show the dynamics of the sensitive and resistant populations, respectively, under the treatment trajectory without error, whereas the corresponding solid lines show these dynamics under the trajectory with error. **A** If the tumor burden almost reaches the progression threshold during the period without treatment, an error can increase the measurement and lead to the declaration of the treatment failure. **B** Measurement error reduces the observed tumor burden even though the true burden has already reached *N*_max_. As a result, treatment is not resumed when it should be, leading to an additional treatment holiday during which the tumor burden exceeds the progression threshold. **C** The top graph shows the treatment dynamics without measurement error, whereas the bottom graph shows the dynamics with a measurement error at the 30-day test point. The measurement error raises the observed tumor burden above *N*_min_, causing treatment to be continued for an additional 30 days. This leads to a resistant population overgrowing the sensitive one, resulting in the loss of one treatment cycle and a reduction in TTP.

Measurement error can lead to a higher TTP in certain cases. For example, if the measured tumor burden is close to, but has not yet reached, the upper bound, measurement error may cause it to exceed the threshold and prematurely trigger treatment resumption. In this case, we avoid leaving a patient for another 30 days without treatment, during which the tumor burden might cross the progression threshold. Similarly, measurement error may prevent an unnecessary treatment interval that would otherwise accelerate resistance development.

In the simulations, measurement error decreased median TTPs for all patients under both Zhang et al.’s and the containment protocol, indicating that the negative effects of measurement errors outweigh any potential benefits as the error magnitude increases. However, measurement errors lead to substantial variability in outcomes for the same patient. Figure 4B shows the distribution of TTP across simulation runs for different values of the log-scale standard deviation *σ* for three patients. Median TTP decreases as *σ* decreases, whereas the interquartile ranges remain wide. Two density peaks-one at higher TTP values and one at lower TTP values-indicate two distinct treatment trajectories: one largely unaffected by measurement error and another strongly compromised by it. For example, for patient 1045, Zhang et al.’s protocol with *σ* = 0.1 can result in a TTP close to the protocol baseline (significantly higher than under MTD) or below the MTD baseline. However, for larger values of *σ*, variability decreases substantially. Most simulation runs yield TTP values below the MTD baseline, thereby eliminating the benefits of Zhang et al.’s protocol.

**Figure 4.**
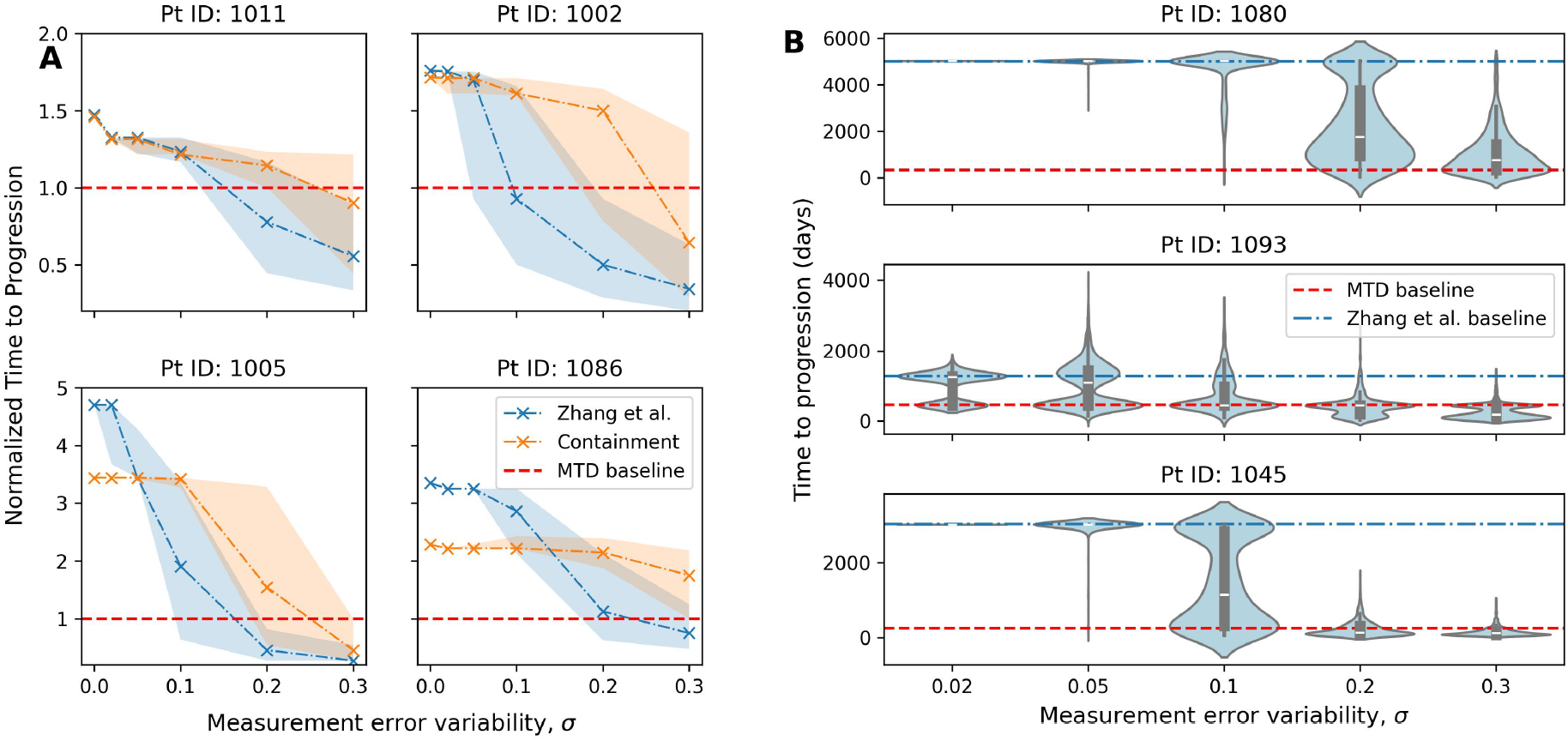
Effects of increasing mean measurement error on the median and distribution of TTP for selected virtual patients. **A.** Zhang et al.’s protocol is more susceptible to measurement error than the containment protocol. The blue line and shaded area show the median TTP and interquartile range for patients under Zhang et al.’s protocol. The orange line and the shaded area show the same for the containment protocol. For the four illustrated patients, Zhang et al.’s protocol yields a higher TTP without measurement error in the absence of measurement error, but median TTP decreases more rapidly than under the containment protocol as measurement error variability *σ* increases. **B**. Higher values of *σ* result in a high variability of TTP across simulations. Violin plots show distributions of TTPs across 1000 simulation runs for each value of *σ*.

We observed that Zhang et al.’s protocol is more sensitive to the measurement error than the containment protocol. While without the measurement error, TTP of Zhang et al.’s protocol is not lower than that of the containment protocol for all virtual patients, with *σ* = 0.3 this is true only for 9 out of 38 patients. In many cases, the decline in median TTP with increasing variance (as determined by *σ*) is steeper under Zhang et al.’s protocol than under the containment protocol (Figure 4A). This is explained by Zhang et al.’s protocol having a higher upper bound than the containment protocol. Therefore, if the true tumor burden exceeds *N*_max_ but, due to measurement error, treatment is not resumed, the tumor burden under Zhang et al.’s protocol is more likely to surpass the progression threshold before the next assessment.

### 3.4 Interaction between error and delay

For each combination of appointment delay *λ* and measurement error *σ*, we performed 1000 simulation runs and obtained the resulting TTP values. Because the distribution of TTP for a given patient under fixed conditions is often non-normal and skewed, we based the analysis on the median TTP for each parameter combination. The heatmaps illustrate changes in median TTP with increasing *σ* and *λ* for each patient 5. Median TTP values are normalized to the protocol baseline, defined as the TTP under the protocol without measurement error or delay. Virtual patients are classified into four clusters using K-Means clustering implemented in the `scikit-learn` Python library (Pedregosa et al., 2011). The clustering is based on a sensitivity decomposition that quantifies the magnitude of the effects of measurement error, appointment delay, and their combination on each patient’s median TTP (Appendix A4). The four clusters, the corresponding sensitivity decompositions, and the normalized median TTP heatmaps for the 38 virtual patients under Zhang et al.’s protocol are shown in Figure 5.

**Figure 5.**
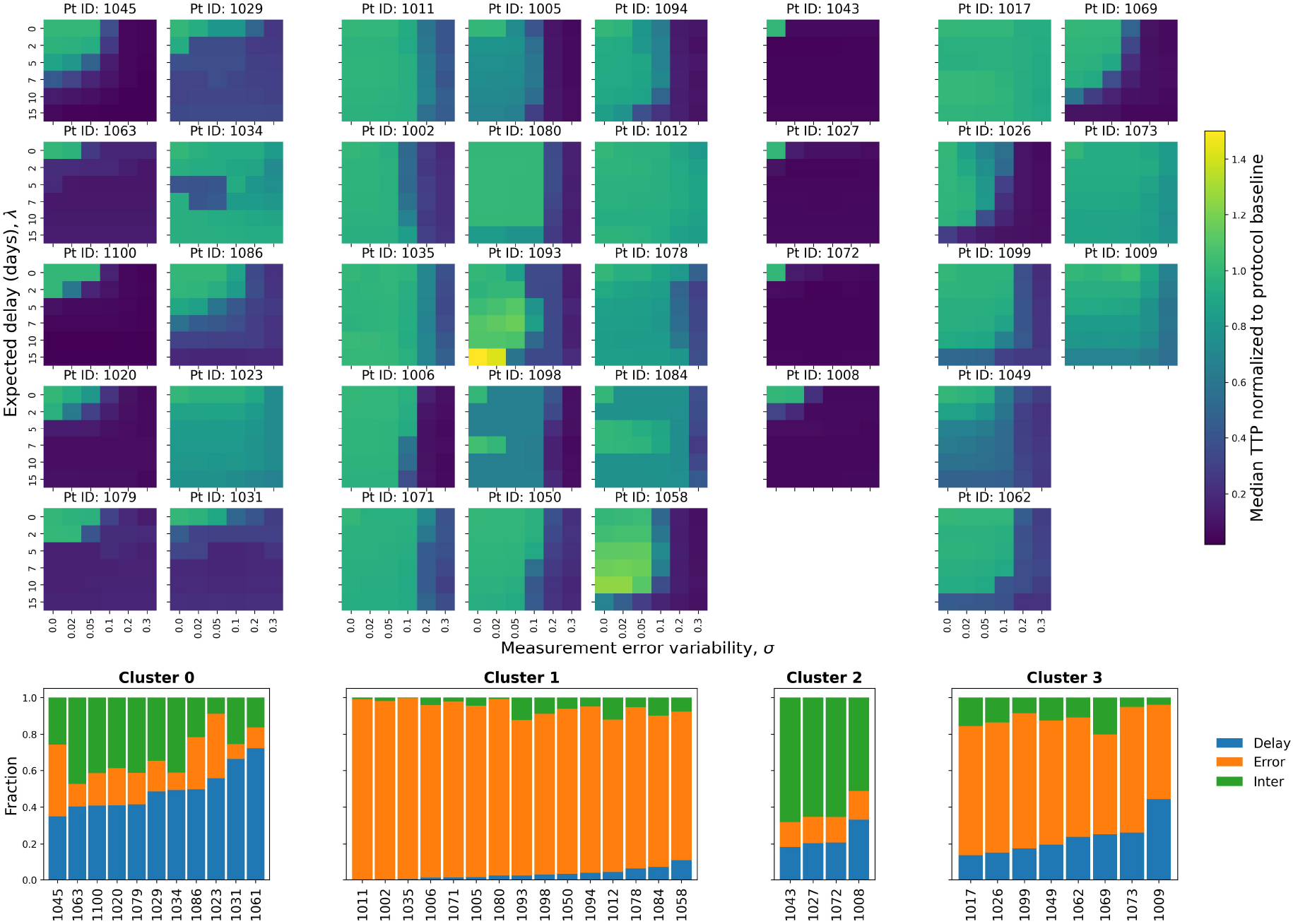
Change in median TTP, normalized to the protocol baseline, under Zhang et al.’s protocol for 38 virtual patients grouped into four clusters based on their sensitivity decomposition of median TTP variation. The heatmaps show the change in median TTP, normalized to Zhang et al.’s baseline (the TTP under no delay, no error conditions), across combinations of expected delay *λ* and error variance determined by *σ*. Each heatmap corresponds to one virtual patient. The virtual patients are divided into four clusters based on their sensitivity decomposition of the median TTP variance. The blue, orange and green bars indicate the fractions of the variation attributed to delay, error and their interaction, respectively.

The largest cluster (cluster 1) contains virtual patients for whom the measurement error plays a major role in the variance of the median TTP. Those patients are almost unaffected by appointment delays, but a measurement error changes their median TTP significantly. In the heatmaps, the differences between columns are more apparent than those between rows. This is especially evident in cases of large values of *σ* = 0.2 or *σ* = 0.3. In contrast, for virtual patients in cluster 0, appointment delay plays a more prominent role; however, its impact remains smaller than the effect of measurement error observed in cluster 1. For many patients in this cluster, delays result in a notable drop in median TTPs. For virtual patients in cluster 2, combination of measurement errors and delays is prevalent in the sensitivity decomposition. The heatmaps of median TTP changes indicate highly sensitive tumor dynamics, for which even small measurement errors or appointment delays lead to premature protocol failure. Finally, cluster 3 consists of patients for whom measurement error plays an important role, as in cluster 1, but appointment delay also contributes. The effects of error and delay reinforce each other. For the same value of *λ*, a higher value of *σ* leads to a lower median TTP, and vice versa. For some patients in this cluster (patients 1017 and 1073), both components affect median TTP, but the magnitude of the effect is moderate. They present a favorable case in which neither error nor delay eliminates the benefits of ECT. Overall, for the majority of patients, the effect of measurement error on median TTP is greater than that of appointment delay.

For the containment protocol, K-means clustering of the median TTP variance decomposition yields the same cluster categories (Figure 6). The heatmaps show the changes in median TTPs, normalized to the patient’s TTP under the containment protocol without error or delay. Compared with Zhang et al.’s protocol, changes in median TTP are less pronounced. We then compare median TTPs under Zhang et al.’s and the containment protocols for each combination of *λ* and *σ* for all patients. Table 1 shows the percentage of patients (out of 38) for whom the median TTP under Zhang et al.’s protocol is not lower than that under the containment protocol for a combination of delay and error parameters.

**Table 1.** The percentage of patients out of 38 with the median TTP under the containment protocol lower than that under Zhang et al.’s protocol across the values of expected appointment delay *λ* and measurement error variance determined by *σ*. The columns indicate the values of log-scaled standard deviation *σ* of measurement error and the rows indicate the mean values of the appointment delay *λ* in days. While the TTP under Zhang et al.’s is not lower than that under the containment protocol for almost all virtual patients with small *σ* and *λ* values, the fraction decreases as these values increase, indicating that the containment protocol is more robust to varying clinical conditions.

| $\lambda$ | $\sigma$ | | | | | |
| --- | --- | --- | --- | --- | --- | --- |
|  | 0.00 | 0.02 | 0.05 | 0.10 | 0.20 | 0.30 |
| 0 | 100 | 97 | 89 | 71 | 34 | 26 |
| 2 | 97 | 97 | 87 | 53 | 34 | 21 |
| 5 | 89 | 92 | 76 | 47 | 37 | 29 |
| 7 | 84 | 84 | 71 | 34 | 34 | 26 |
| 10 | 87 | 79 | 79 | 53 | 42 | 32 |
| 15 | 84 | 87 | 71 | 50 | 45 | 34 |

**Figure 6.**
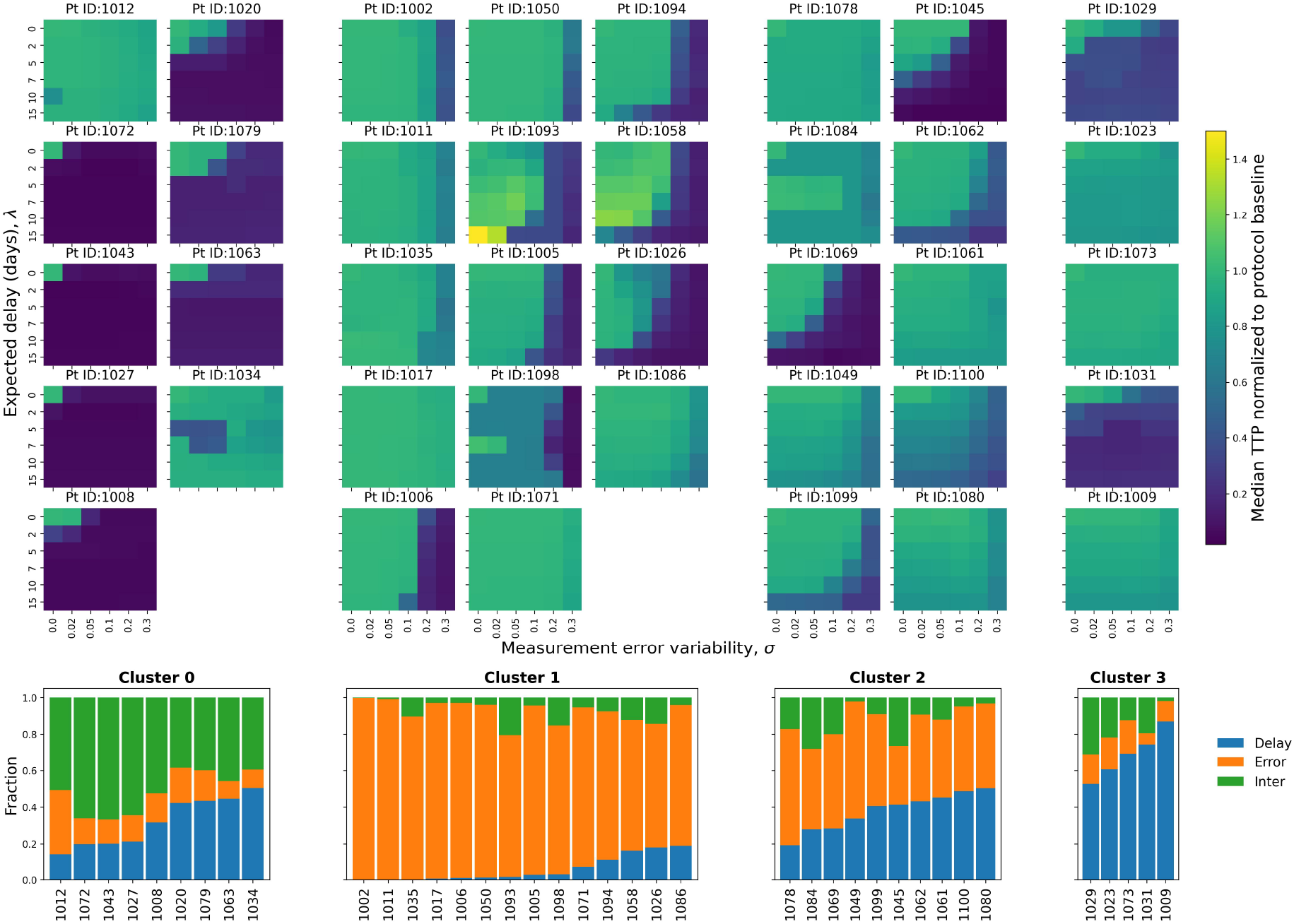
Change in median TTP of virtual patients under the containment protocol across delay and measurement error combinations, grouped into four clusters based on the fraction of variation attributed to delay, measurement error and their interaction. The heatmaps show the change in median TTP under the containment protocol with increasing expected delay *λ* and error variance determined by *σ*. Each heatmap corresponds to one virtual patient. Median TTP values are normalized to the TTP under the containment protocol in the absence of delay and measurement error for each patient. The virtual patients are divided into four clusters based on the decomposition of median TTP variance. The blue, orange and green bars indicate the fractions of the variation attributed to delay, error and their interaction.

A significant decrease in median TTP with an increase of *λ* and *σ* does not necessarily imply that the median TTP drops below that under MTD. As TTP under Zhang et al.’s protocol is higher than that under the containment protocol without error or delay for the majority of patients, a more apparent relative decrease in median TTP does not necessarily indicate the protocol’s failure for higher values of *λ* and *σ*. To assess how probable the premature failure of the evolutionary therapy protocols is with an increase in error variability and expected delay, we compared the fraction of simulation runs out of 1000 that resulted in TTP exceeding that under MTD for each combination of *λ* and *σ*. Figure 7 compares the number of simulation runs that outperform MTD for the two evolutionary protocols for an example patient 1069. Low values of *λ* and *σ* have little effect on TTP of evolutionary protocols, whereas their higher values lead to failure in the majority of cases. For most parameter combinations, the containment protocol yields a larger number of simulation runs with TTP exceeding the MTD baseline than Zhang et al.’s protocol.

**Figure 7.**
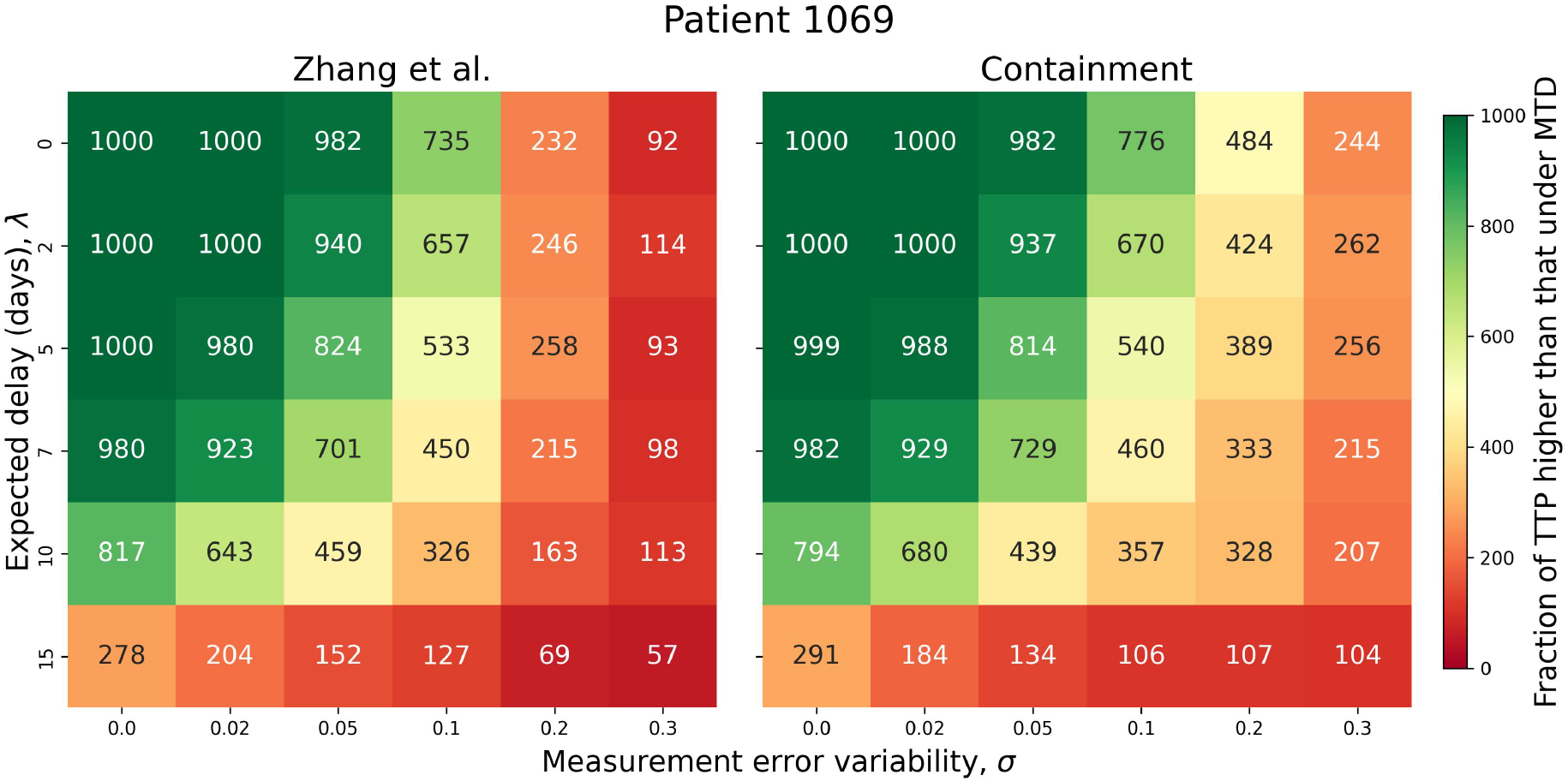
Number of simulation runs (out of 1000) across combinations of *λ* and *σ* that resulted in TTP exceeding the MTD baseline for patient 1069 under Zhang et al.’s and the containment protocols. In general, the containment protocol has a higher fraction of simulations that outperform MTD across the simulated conditions.

To investigate the role of individual tumor growth rates in determining sensitivity of the cancer dynamics to delay and measurement error, we simulated a 30-day unmedicated tumor growth period at the start of therapy and calculated the corresponding linear growth rates for all patients (Figure A7). Figure A5 shows the changes in the fraction of simulations with TTP higher than under MTD across combinations of *λ* and *σ* for all 38 virtual patients, sorted by increasing linear unmedicated growth rate. Virtual patients with lower growth rates are generally less affected by appointment delay and measurement error; however, growth rate alone does not explain all observed variability in the results (Figure A5 and Figure A6).

### 3.5 Dynamically adjusted containment level

We implemented a dynamically adjusted containment protocol that estimates the untreated linear tumor growth rate and determines the maximum containment level that prevents the tumor burden from crossing the progression threshold between scheduled assessments. Because linear estimation of the growth rate can be imprecise, the dynamic containment protocol led to premature failure in 5 patients even in the absence of measurement error or appointment delay. In the remaining 33 out of 38 patients, the dynamic containment resulted in a TTP not lower than the standard containment protocol. In 13 of these patients, the resulting TTP exceeded that achieved under the other evolutionary protocols, and in 19 patients it matched that of Zhang et al.’s protocol, which is never lower than the containment protocol in the absence of delay and measurement error.

The higher TTP under the dynamic containment protocol is attributable to the higher containment level that it seeks to maintain. However, there is an increased risk of premature failure. Firstly, the dynamic containment protocol sets the containment bound as high as possible, but so that the tumor size does not exceed the progression threshold between tests. The linear growth rate used for that might not be a sufficiently accurate estimate and could overestimate a safe containment level. Second, the calculations assume test intervals without delay. Therefore, the protocol is sensitive to unexpected appointment delays. Lastly, the growth rate is estimated from tumor burden measurements, which may contain errors. If the previous measurement is overestimated and the subsequent one underestimated, the estimated growth rate may be substantially lower than the true rate, leading to a containment bound that is set dangerously high. In this case, measurement error affects not only treatment decisions at individual assessment points but also the design of the protocol itself.

To assess the safety of the dynamic containment protocol in clinically realistic conditions, we compared how the fraction of simulation runs with TTP higher than that under MTD changes with increasing values of *λ* and *σ*, and compared it with constant-schedule evolutionary therapy protocols.

Figure 8 shows the effect of measurement error and appointment delay on the fraction of TTPs higher than under MTD of the 7 patients under the three evolutionary therapy protocols. The heatmaps show the number of simulation runs (out of 1000) that resulted in TTP higher than the MTD baseline, indicating the success of the evolutionary therapy protocol. Patients are ordered according to increasing linear growth rate of the untreated tumor over a 30-day period (Figure A7). The virtual patient with the lowest growth rate has a low risk of premature failure, which occurs only for large measurement errors corresponding to high values of *σ*. The higher linear growth rate corresponds to a larger effect of error and delay, with the last patient being affected even by small *λ* and *σ*. We can observe that the dynamic protocol has a rapidly increasing number of simulation runs resulting in premature failure with increased measurement error and appointment delay. This is especially noticeable in patients 1071 and 1069. In general, the dynamic containment protocol is more sensitive to measurement errors and appointment delays than constant protocols.

**Figure 8.**
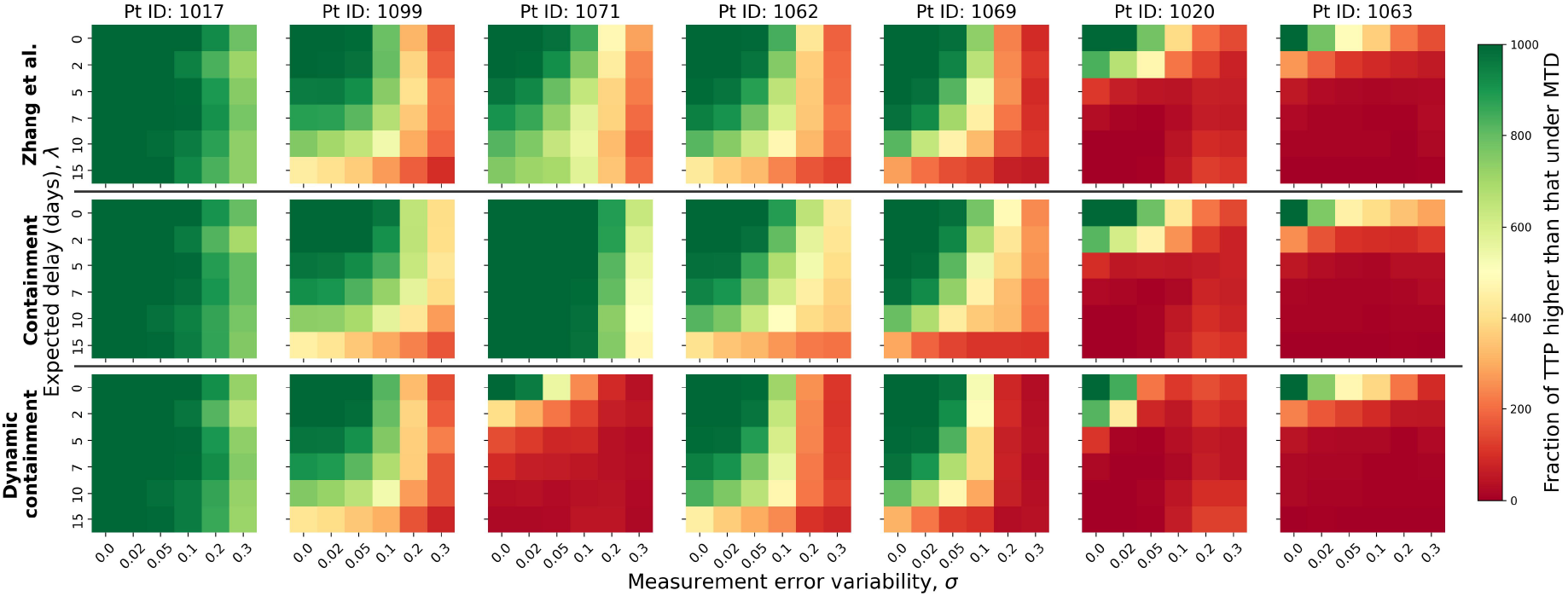
Comparison of Zhang et al.’s, containment and dynamic containment protocols under increasing expected appointment delay (*λ*) and measurement error variability (*σ*). The heatmaps show the fraction of 1000 simulation runs resulting in TTP higher than that under MTD across combinations of *λ* and *σ* for 7 patients under the three treatment protocols. In general, the dynamic containment protocol is especially susceptible to errors and delays, while the containment protocol is the most robust.

## 4 Discussion

In this study, we simulated clinically realistic conditions, including monthly assessment intervals, appointment delays, and measurement errors, and evaluated their effects on the outcomes of evolutionary cancer therapy protocols. We demonstrated that longer appointment delays and larger measurement errors can undermine the benefits of evolutionary therapy. However, these adverse effects can be partially mitigated through appropriate treatment protocol design. Based on these findings, we aim to facilitate the dialogue between physicians and evolutionary therapy modelers by projecting modelbased predictions of tumor dynamics onto more realistic clinical conditions and thereby supporting the design of future clinical trials.

A particularly important risk to consider is premature failure of ECT protocols driven by rapid tumor growth during treatment holidays. In the simulations, progression was defined as the tumor burden exceeding its initial size by 20% regardless of whether the medication is on or off at the time. It differs from the definition of progression used in an ongoing clinical trial on prostate cancer, where the progression can only be declared if the treatment is on. Using the latter definition would improve simulation outcomes in cases where the tumor burden slightly exceeds the progression threshold, but the sensitive population is still present, and the next treatment cycle is possible. However, allowing tumor burden to exceed the progression threshold remains undesirable, as it may exacerbate cancerrelated symptoms and increase the likelihood of additional tumor mutations (He et al., 2025; West et al., 2023). A subset of simulated patients was predicted to progress during the first treatment cycle due to the rapid growth of the untreated tumor. Identifying such cases prior to treatment initiation is challenging, as patient-specific tumor dynamics are typically unknown at baseline. This underscores the need for particularly frequent and careful monitoring during the first treatment cycle. An alternative approach would be to perform an additional assessment on the day treatment is initiated. Because this time point is usually more than four weeks after the baseline diagnostic test, it would enable estimation of the untreated tumor growth rate, a parameter that is critical for the safe implementation of ECT protocols.

Appointment delay affects treatment outcomes by prolonging either the treatment-free interval, which may lead to premature failure, or the on-medication interval, which can promote the growth of the resistant population. If a delay during the treatment-free interval does not result in premature failure, it may be considered beneficial within the modeling framework, as it permits a higher containment level. In models incorporating a cost of resistance, this increases the relative fitness of the sensitive population and thereby prolongs treatment effectiveness. However, in clinical practice, a higher tumor burden would likely mean more discomfort and more prevalent cancer symptoms for the patient, a strong motivation to avoid delays during the off-medication period.

While an appointment delay only postpones a treatment decision, a measurement error can change the decision itself. This leads to an unnecessary interval (30 days in our simulations) with either medication administered, in which case it promotes resistance development, or without medication, in which case there is a risk of the tumor growing too large, leading to premature treatment failure or unwanted symptoms for the patient.

In simulation scenarios with both delay and error, the two factors often amplify each other. A higher error with the same mean delay would result in both a lower median TTP and a higher occurrence of premature failure, and vice versa. We observed that the error has a larger effect on decreasing the median TTP and increasing occurrences of premature failure. Furthermore, in clinical practice, measurement error is less controllable than appointment delay. When a delay occurs, its duration is known and can be incorporated into treatment design. In contrast, measurement error affects the observed tumor burden without revealing the true value, and some level of uncertainty must always be assumed. In our simulations, small values of error variability (*σ* ≤ 5%) had only a minor impact on evolutionary therapy outcomes for the majority of virtual patients. Therefore, reducing measurement error may facilitate the clinical implementation of evolutionary therapy. Technological advances, such as computer vision tools, can improve accuracy and reduce the time required for lung lesion volume estimation (Xie et al., 2021; Primakov et al., 2022). It is also important to consider that measurement error tends to be greater in smaller lesions, which are more difficult to quantify reliably on imaging, leading to increased uncertainty and inter-observer variability among radiologists (Oxnard et al., 2011a; McNitt-Gray et al., 2015; Goodman et al., 2006; Henschke et al., 2016). At the same time, other differences in the response of larger tumors versus smaller tumors to ECT can be present (Gallaher et al., 2022), highlighting the importance of the individual cancer composition in therapy design.

Zhang et al.’s protocol results in a higher TTP than the containment protocol in all patients under no-delay, no-error conditions, due to a higher containment level of the containment protocol. However, the median TTP decreases sharply once errors and delays are introduced. The reason is that a higher upper bound in combination with appointment delays and measurement errors leads to a high risk of premature failure. To partially mitigate this risk, we implemented a dynamically adjusted test intervals (Appendix A5). The idea is to adapt the intervals between assessments based on the observed tumor decrease and regrowth rates, in order to better maintain tumor burden within the bounds of Zhang et al.’s protocol. In the case of insignificant error and delay, this approach performs worse than standard Zhang et al.’s protocol, as the resulting average containment level is lower, but it is more robust to higher errors and longer delays and might be preferred in clinical practice because it provides greater control over tumor size. The containment protocol, however, was demonstrated to be “safer” than Zhang et al.’s protocol even with a dynamic schedule. In the simulations, the containment protocol resulted in a premature failure less often across all combinations of measurement error and appointment delay. A protocol with a single containment bound is more robust to real clinical conditions of rather long intervals between tests. With this protocol, the tumor burden cycles around the bound and can grow or shrink further from it. As the gap between the measured tumor burden and the protocol threshold increases, the risk that measurement error will shift the observation across the bound and alter the treatment decision decreases.

We also tested the dynamically adjusted containment protocol, in which the containment bound is determined based on the linear estimate of the tumor growth rate during the medication-free interval. As the protocol aims to set the containment level as high as possible, it led to a higher TTP than that under Zhang et al.’s and the containment protocols in 19 out of 38 patients in the no-delay, no-error setting. However, with a large error and long delay, this protocol performs worse than the constant-schedule protocols in terms of TTP. The estimated containment level is highly affected by measurement errors, because the growth rate is calculated from the erroneous measurements, and by appointment delays, because the tumor burden is supposed to grow to progression lines between tests, but does not account for unexpected delays. As a result, an imprecise growth rate estimate may lead to an excessively high containment level and, consequently, premature failure.

There is a trade-off between achieving greater TTP benefits by maintaining a higher tumor burden and enhancing safety by keeping the burden lower. When dealing with a fast-growing type of cancer and accounting for clinically realistic intervals between tests, avoiding tumors going untreated for a long time should be a priority. As individual cancer dynamics play a crucial role in ECT outcomes, we recommend starting with a conservative threshold of 50% of the initial tumor burden. For patients with a high rate of tumor regrowth, careful follow-up should be a priority. This can include high-resolution scans to reduce measurement error, more strategic appointment planning and avoiding delays. If the rate of regrowth is moderate, the focus can be on maximizing TTP benefit by setting the containment level higher. However, it is important to recognize the risks of such an approach, as it is particularly sensitive to errors and unexpected delays.

The main operational challenge in implementing ECT for metastatic NSCLC is the lack of a universally validated serum biomarker that consistently tracks tumor burden across patients (Hendriks et al., 2023). The assessment of tumor burden in clinical practice relies predominantly on radiological imaging (Eisenhauer et al., 2009; Schwartz et al., 2016), which is limited both in precision and in the frequency at which it can be performed. Circulating tumor DNA (ctDNA) represents a promising emerging biomarker in NSCLC. Recent prospective real-world data in KRASG12C-mutated NSCLC demonstrate that early changes in ctDNA variant allele frequency correlate with progression-free and overall survival and may precede radiographic progression (Ernst et al., 2024). These findings suggest that dynamic molecular monitoring could enable earlier detection of treatment response or failure compared to imaging alone. Although routine implementation of ctDNA for high-frequency evolutionary treatment guidance remains under development, continued advances in liquid biopsy technologies may facilitate more precise and frequent tracking of tumor dynamics, thereby supporting the future clinical implementation of evolutionary therapy in NSCLC.

The proposed modeling framework provides an opportunity to evaluate potential clinical scenarios of evolutionary cancer therapy for NSCLC and assess its clinical feasibility before real-world implementation and resource allocation. Nevertheless, several limitations of the modeling approach should be considered. First, the study relies on virtual patients based on clinical cancer dynamics observed under continuous MTD treatment. The cancer dynamics under evolutionary therapy protocols were then predicted by the model and may not fully reflect those of real patients. Further, the resistant cell population was assumed to be almost absent at the beginning of the treatment, based on clinicians’ observations that patients rarely have preexisting resistance to the first-line osimertinib treatment. A larger initial resistant population would likely change the predicted evolutionary dynamics and treatment outcomes.

The linear growth rate estimation used in the dynamic containment protocol and for dynamic test scheduling represents a simplified approximation of tumor dynamics. Incorporating a cancer growth model can more accurately suggest an optimal containment threshold. However, in practice, it is not clear which of the various cancer growth models to use or how to properly parameterize them with often scarce medical data. Further, such a modeling task would likely require a multidisciplinary team involved in designing a patient’s treatment trajectory, which is often not feasible in wide clinical practice. Linear growth cannot be safely applied to high-precision protocols and scheduling tailoring, but if the aim is simply to identify slow and fast growers, it may be sufficient. Finally, we only considered binary evolutionary therapy protocols, in which the treatment is either on or off at each time point, and not dose-modulation ECT protocols (Cunningham et al., 2020a; Hockings et al., 2023; Salvioli et al., 2024). Such protocols may be a preferable option for rapidly growing cancers, where complete treatment interruption for a prolonged period may be undesirable. However, the dose-modulation protocols would likely require even more accurate tracking of disease progression and more frequent follow-up, potentially including the development of biomarkers. The feasibility of such treatment protocols in clinical realities should be evaluated in future studies.

This study bridges evolutionary cancer therapy and clinical reality by incorporating operational constraints into therapy design. Simulation-based insights can inform future clinical trials and facilitate the translation of ECT into metastatic NSCLC care.

## APPENDIX A A1 Parameterizing the cancer dynamics model with patient data and creating virtual patients

In the study, we used data from the START-TKI trial (NCT05221372). The START-TKI trial is conducted in accordance with the Declaration of Helsinki. Ethical approval for the trial was granted by the Medical Ethics Review Committee (METC) of Erasmus Medical Center, Rotterdam, The Netherlands (MEC 16-643) in accordance with the Dutch Medical Research Involving Human Subjects Act (WMO). Informed consent was obtained from all patients prior to enrollment in the trial. This study does not use any identifiable or individual-level patient data from START-TKI trial.

All patients are patients with metastatic non-small cell lung cancer treated with osimertinib. The data consist of time-series measurements of the tumor’s longest diameters across three projections. We defined tumor volumes as volumes of ellipsoids with radii equal to the observed measurements. We calculated the total tumor burden as the sum of all measured lesions at every time point. If measurements of different lesions were performed within a week of each other, we assumed they were taken on the day of the primary lung lesion measurement. As we are interested in tumor response to osimertinib treatment, tumor measurements obtained after switching to a different medication were excluded.

The polymorphic Gompertzian model was fitted to the time-series data for total tumor volume for 37 patients. As the patients were treated with first-line osimertinib, we assumed almost no preexisting resistance and set the initial total tumor burden to be *N* (0) = *S*(0) + *R*(0), with the initial resistant and sensitive populations set to *R*(0) = 5 ·10^−6^ *N* · (0) and *S*(0) = *N* (0) − *R*(0). The remaining model parameters were estimated using Python 3.12.3 with the differential evolution algorithm (Storn & Price, 1997) implemented in the `scipy.optimize` library. Differential evolution (DE) is a global optimization algorithm designed for continuous, non-linear problems. The model was fitted to the data by solving the differential equations with the Runge–Kutte 45 method from the `solve_ivp` module in the `scipy.integrate` library.

In the fitting procedure, we applied the following bounds for the parameter values. For the growth rates *ρ*_*R*_ and *ρ*_*S*_ the bounds are 0.0001 and 0.06. We set treatment sensitivity *λ* to be between 0.0001 and 0.2.The lower and upper bounds for the competition coefficients *α*_SR_ and *α*_RS_ were set at 0.1 and 10, respectively. The bounds for the carrying capacity *K* were set individually for each patient as 1.3*N* (0) and 5 ∗ 10^5^. The parameter bounds and the fitting procedure are analogous to the previous study on similar patient data (Jansén-Storbacka et al., 2026).

Due to imperfections in the data and potentially limitations of the fitting procedure, some of the obtained parameter values were on the bounds, resulting in unrealistic cancer dynamics under treatment. Some patient records showed no tumor regrowth because the patients were still in treatment, or only slight regrowth, which, however, was sufficient to declare progression under RECIST criteria. According to RECIST, progression is declared when there is a 20% increase in one of the longest diameters from its minimum size (Eisenhauer et al., 2009). In the total tumor volume, such an increase can be very small and may be ignored during model fitting. In these cases, the model estimates that resistance develops very slowly, resulting in TTP above 3000 days. Another abnormality in the data is the extremely rapid tumor growth without treatment, which likely stems from the model’s growth rate values being too high. As the model is fitted to tumor dynamics under continuous treatment, extreme growth rate values are compensated by the treatment variable and only become apparent in the protocols with treatment holidays.

To avoid these abnormalities and broaden the range of potential dynamics to consider, we created virtual patients by sampling parameters from a multivariate normal distribution derived from the fitted models. We applied the same parameter bounds that were used in the fitting procedure. To ensure clinical realism, we only sampled patients with TTP below 3000 days under continuous treatment.

## A2 Choosing the optimal protocols’ bounds

For Zhang et al.’s protocol, we tested various values of *N*_max_ = *k*_max_*N* (0) with *k*_max_ ∈ {0.6, 0.7, 0.8, 0.9, 1.0}). For the containment protocol, we tested containment levels *N*_cont_ = *k*_cont_*N* (0) with *k*_cont_ ∈ {0.5, 0.6, 0.7, 0.8, 0.9, 1.0}. For each level, we simulated treatment trajectories for all 100 virtual patients and assessed TTPs and the percentage of premature failures (cases in which the tumor burden exceeds the progression threshold during the treatment-free interval). Table A2 presents the number of cases when the given threshold value leads to the maximum TTP across all *N*_max_ and the percentage of premature failure for each of the threshold values. We considered both indicators to determine the optimal thresholds. For the containment protocol *N*_cont_ = 0.5 has the highest percentage of the best results and the lowest percentage of the premature failures. We therefore decided to focus on this value in the main analysis of the study. In Zhang et al.’s protocol, *N*_max_ = 0.7 has the second highest percentage of the best TTPs (only 1 percent lower than the maximum) and a significantly lower percentage of premature failures than *N*_max_ = 0.8. This level was chosen as optimal for Zhang et al.’s protocol for further simulation.

| Zhang et al. |  |  | Containment |  |  |
| --- | --- | --- | --- | --- | --- |
| $N_{\max}$ | % best results | % premature failure | $N_{\text{cont}}$ | % best results | % premature failure |
| — | — | — | 0.5 | 67 | 43 |
| 0.6 | 75 | 44 | 0.6 | 57 | 51 |
| 0.7 | 79 | 45 | 0.7 | 60 | 55 |
| 0.8 | 80 | 53 | 0.8 | 58 | 63 |
| 0.9 | 79 | 63 | 0.9 | 58 | 71 |
| 1.0 | 69 | 74 | 1.0 | 51 | 82 |

## A3 Increase of TTP due to appointment delay

In a few particular cases, appointment delay can increase TTP under Zhang et al.’s protocol. Three patients who demonstrated premature failure under Zhang et al.’s protocol but not under the containment protocol showed an increase in median TTP with increasing mean appointment delay (Figure A1). This reflects the disadvantage of the strict upper bound in Zhang et al.’s protocol. If the current measurement is close to the upper bound but has not yet reached it, the treatment pause continues for an additional 30 days until the next test, during which the tumor burden grows to the progression level. In this case, a one-week delay of the appointment would lead to the restart of the treatment at the current appointment, thereby prolonging or avoiding premature failure (Figure A1).

## A4 Sensitivity decomposition of the median TTP values

For each virtual patient, we obtained a 6-by-6 matrix of median TTPs *Y* ∈ ℝ^6*×*6^ as the results of 1000 simulation runs. The rows and columns of the matrix correspond to the mean appointment delay *λ* and measurement error *σ*, respectively. For each patient and each treatment protocol (Zhang et al.’s and containment), we quantified the variability in median TTPs arising from delay, error, and their combination. For this, we implemented the following procedure, based on the two-factor ANOVA variance decomposition. A full parametric ANOVA was not performed because the simulated outcomes do not meet the normality assumption required for the test. Our focus in this analysis was on variance attribution to delay and error rather than on hypothesis testing.

We first calculate the overall mean across all delay-error combinations as

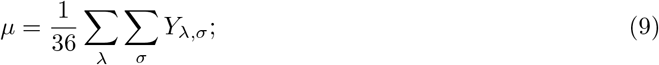

and then the total variance as the mean squared deviation from the overall mean:

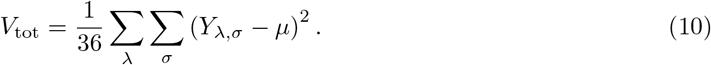

We then compute the variance attributed to delay as:

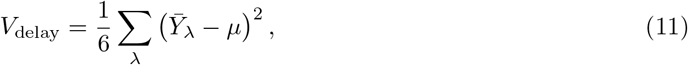

where 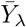 is the average over measurement error for each mean delay value:

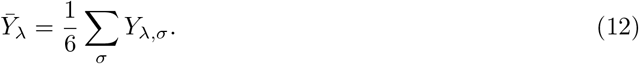

Similarly, the variance attributable to measurement error is

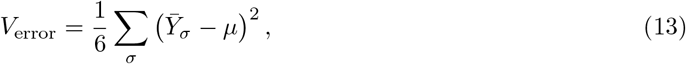

where 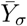 is the average over delay for each mean measurement error value:

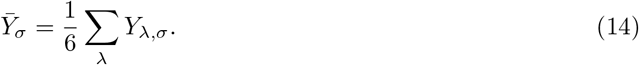

We then normalized the variances attributed to the delay and error to the total variance:

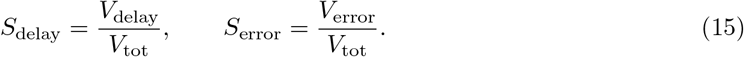

The remaining proportion of variance was assigned to the interaction of delay and error:

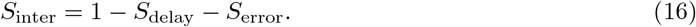

For each patient and each protocol, we calculated what fraction of the variance is attributed to delay, error and their combination (*S*_delay_, *S*_error_, *S*_inter_). Based on these values, the virtual patients were clustered into 4 clusters using K-Means clustering implemented in `scikit-learn` Python library (Pedregosa et al., 2011).

## A5 Zhang et al.’s protocol with dynamic test schedule

We tested the scenario where the time of the next follow-up under Zhang et al.’s protocol is determined based on the rate of tumor burden progression. The dynamic scheduling uses linear estimation of tumor growth without medication and tumor decline under treatment to determine how long it would take for the tumor burden to grow to *N*_max_ or to shrink to *N*_min_. Based on this information, the test intervals are updated. In clinical practice, an interval of less than a month is unlikely to be feasible. At the same time, not performing a follow-up for longer than 3 months is dangerous for cancer patients. We therefore enforced the intervals calculated from the linear growth/decline rate to be between 30 and 90 days.

We implemented rounding up in the dynamic scheduling. If at the test point it is observed that the tumor burden decreased under medication but has not yet reached *N*_min_, the time needed to reach the threshold is calculated based on the tumor burden’s linear decline rate. If it would require less than 14 days, roughly half the standard test interval, the treatment is resumed. Similarly, if the tumor burden at the end of a treatment-free interval has not reached *N*_max_ but is estimated to do so within 14 days, the treatment is resumed. This allows to avoid the situations where the tumor burden almost reaches *N*_min_ or *N*_max_, but has to go another 30 days with or without medication, respectively. That can lead to either overtreating or a higher risk of premature failure and symptoms related to higher tumor burden. From a medical perspective, it is undesirable to allow the tumor burden to reach the initial size or exceed it. Optimizing test scheduling helps maintain tumor burden within the bounds set by Zhang et al.’s protocol, thereby improving therapy outcomes for some patients. At the same time, for some patients, the required test frequency under dynamic scheduling would be lower than every 30 days, thereby decreasing the cost of treatment and the burden on healthcare facilities.

### A5.1 Dynamic test scheduling algorithm

Let *t*_0_ = *t*(0) *< t*_1_ *<* … *< t*_*K*_ = min *t* : *N* (*t*) *> N*_prog_ denote the discrete testing time points at which tumor burden is assessed, repeated until progression, and *N*_mes_(*t*_*i*_) the corresponding tumor burden. The treatment status at time *t* is denoted by *C*(*t*) ∈ {0, 1}, where *C*(*t*) = 1 indicates treatment on and *C*(*t*) = 0 treatment off. At each measurement time *t*_*i*_, we estimate the linear rate of change of tumor burden over the previous interval based on the medication on/off status.

#### Rate estimation

Let *τ*_*i*_ be the most recent time prior to *t*_*i*_ at which the treatment status was switched, *τ*_*i*_ = max {*t*_*j*_ *< t*_*i*_ : *C*(*t*_*j*_) ≠ *C*(*t*_*i*_)}, and *N*_mes_(*τ*_*i*_) the tumor burden at that time.

- If *C*(*t*_*i*−1_) = 1 (treatment on) and *N*_mes_(*t*_*i*_) *< N*_mes_(*t*_*i*−1_), the decline rate is estimated as

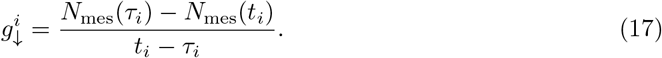
- If *C*(*t*_*i*−1_) = 0 (treatment off) and *N*_mes_(*t*_*i*_) *> N*_mes_(*t*_*i*−1_), the growth rate is estimated as

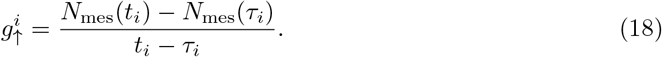

#### Prediction of the bound crossing

- If *C*(*t*_*i*−1_) = 1 and *N*_mes_(*t*_*i*_) *> N*_min_, the predicted time until reaching *N*_min_ is

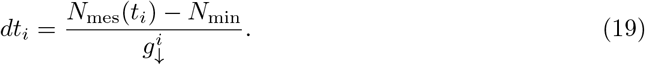
- If *C*(*t*_*i*−1_) = 0 and *N*_mes_(*t*_*i*_) *< N*_max_, the predicted time until reaching *N*_max_ is

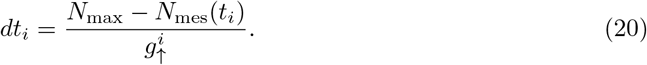

#### Treatment switching and scheduling

If the predicted time to reach the bound satisfies *dt*_*i*_ *<* 14 days, the treatment is switched at *t*_*i*_, i.e.

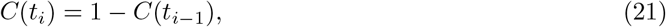

and the next appointment is scheduled based on the rate corresponding to the new regime.

If *dt*_*i*_ ≥ 14 days, the time to the next appointment is set as

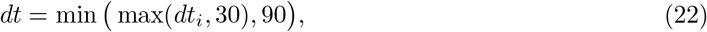

ensuring that the next test occurs no earlier than 30 days and no later than 90 days.

### A5.2 Results

When simulated in no error, no delay conditions, Zhang et al.’s protocol with dynamic scheduling outperforms MTD in terms of TTP for all 38 patients. Further, for 30 patients, the TTP is not lower than that of the containment protocol, and for 10 is higher than that of Zhang et al.’s protocol with a constant schedule. Figure A2 shows an example of virtual patient 1079, for whom the dynamically adjusted test schedule led to an increase in TTP under Zhang et al.’s protocol. The medication periods were adjusted to a single longer interval rather than two standard intervals, resulting in less time on medication and, thus, postponed development of resistance.

We applied appointment delay and measurement error, as previously done for the protocols with constant schedules. For each delay-error combination, we compared the median TTPs between Zhang et al.’s protocol with the dynamic schedule and the constant schedule. Table A1 shows the percentage of virtual patients for whom the median TTP was higher under Zhang et al.’s protocol with the constant schedule than with the dynamic schedule (left) and for whom the median TTP under the containment protocol was higher than under Zhang et al.’s protocol with dynamic schedule (right) for each combination of *λ* and *σ*.

When the measurement error variability, denoted by *σ*, is small, Zhang et al.’s protocol with the constant schedule results in a higher median TTP than Zhang et al.’s protocol with the dynamic schedule for the majority of virtual patients. This is explained by longer treatment-free intervals allowing the tumor burden to grow to a larger size, thereby increasing TTP in the model’s predictions. However, with large values of *σ*, the median TTPs for most virtual patients are higher under the dynamic-schedule Zhang et al.’s protocol. At the same time, the percentage of virtual patients for whom the containment protocol results in a higher median TTP than the dynamic-schedule Zhang et al.’s protocol increases significantly with higher values of *σ*. This indicates that when a large measurement error is present, the containment protocol performs better than Zhang et al.’s protocol, even with a dynamically adjusted schedule.

---

**Figure A1.**
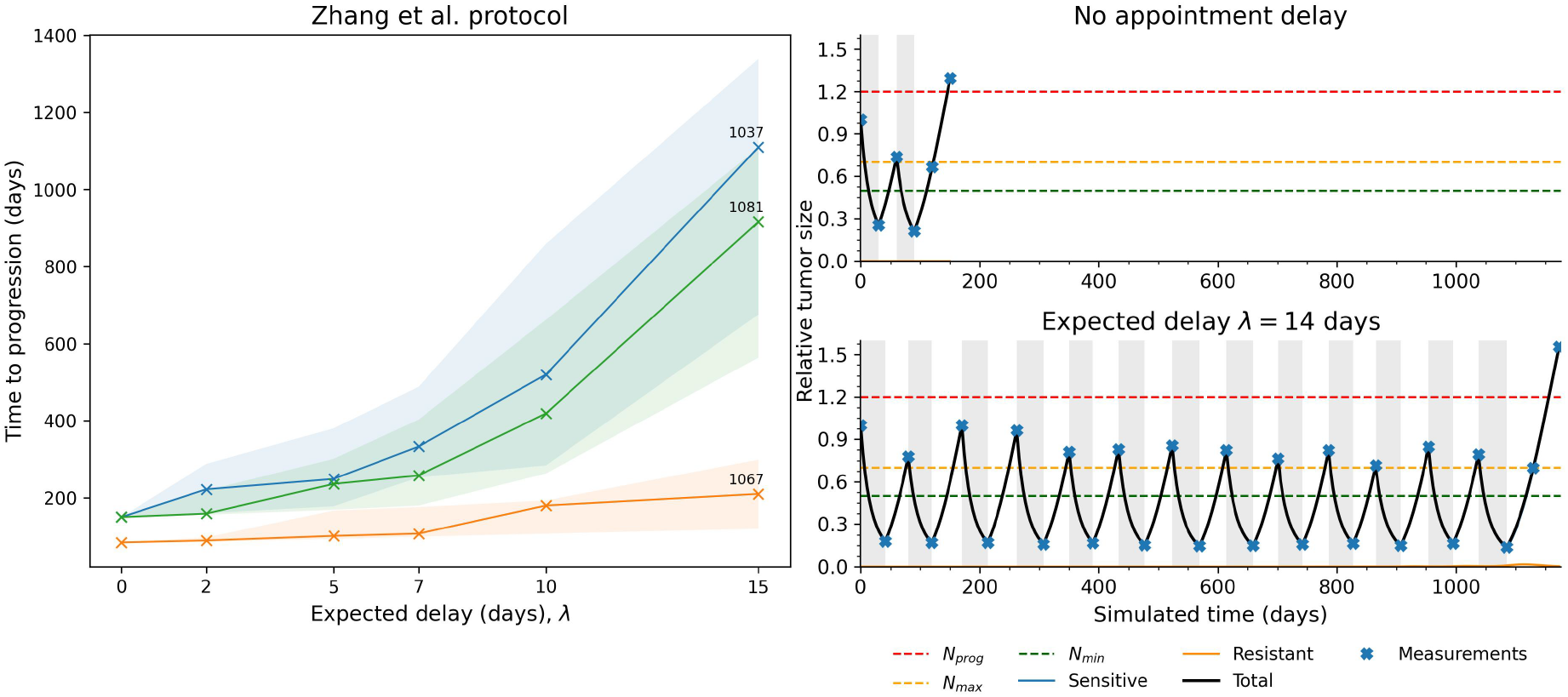
Appointment delay can help to avoid the premature failure of Zhang et al.’s protocol. The graph on the left shows the dynamics of the median TTP for three virtual patients for whom Zhang et al.’s protocol, but not the containment protocol, resulted in premature failure in the delay-free conditions. The shaded area reflects the interquartile ranges. Median TTP increases with the appointment delay for these patients. The right side shows the treatment dynamic of patient 1081 under constant 30 days test intervals (top) and one of the simulation runs with the mean appointment delay of 15 days (bottom). In the first case, the cancer burden does not reach *N*_max_ after 30 days without medication, leading to the continuation of the treatment pause for another 30 days, during which it exceeds the progression threshold. With the appointment delay, the initial interval without treatment is longer than 30 days, allowing the cancer burden to grow beyond *N*_max_ but not beyond *N*_prog_. The treatment is renewed at this time point, resulting in a delay of premature failure and higher TTP.

**Figure A2.**
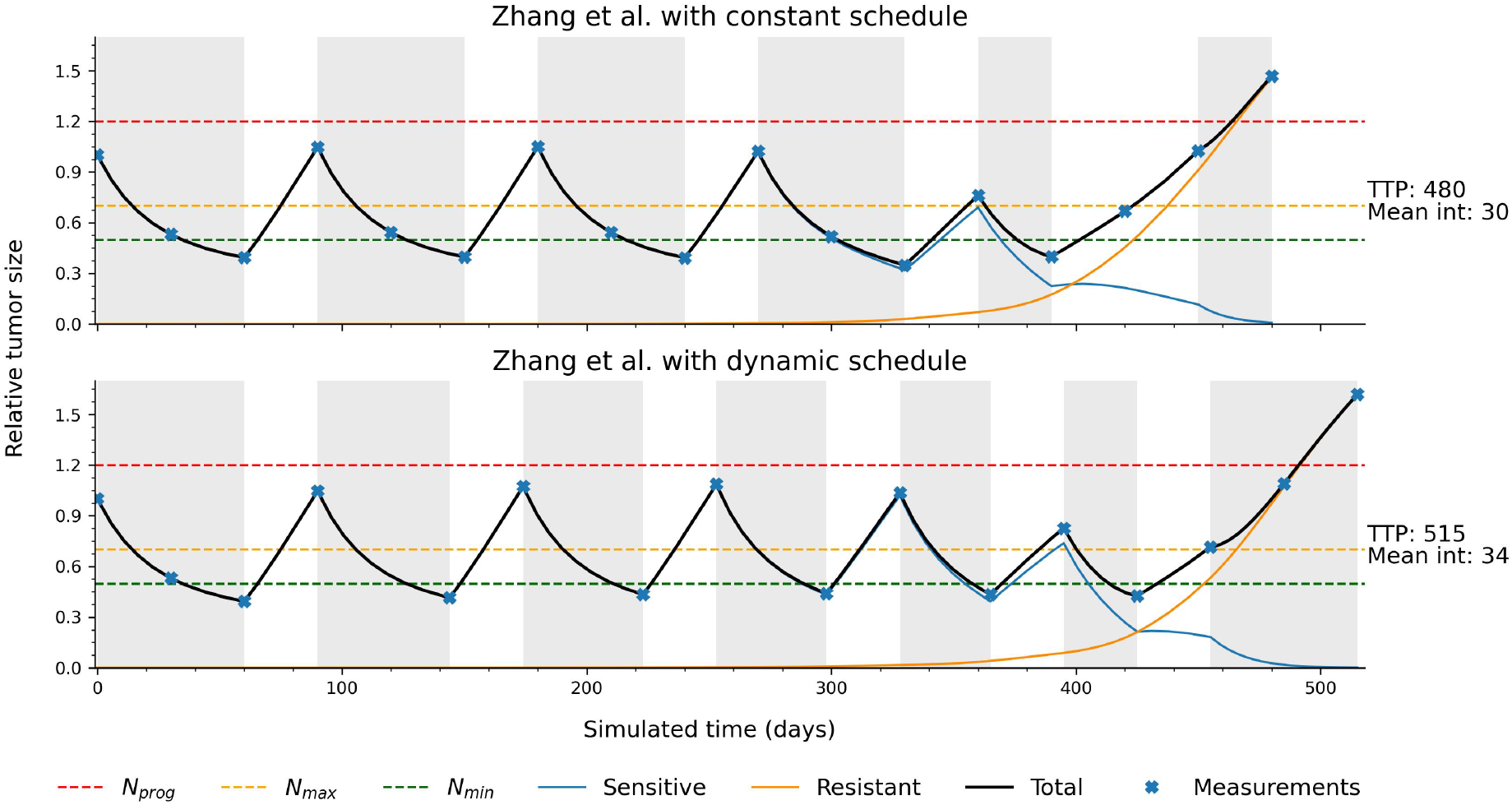
The treatment trajectory of an example patient 1079 under Zhang et al.’s protocol with the constant test schedule (top) and dynamically adjusted test schedule (bottom). Red, yellow, and green dashed horizontal lines indicate the progression threshold, upper and lower bounds of Zhang et al.’s protocol, respectively. The black line shows the total tumor burden, and the blue and orange lines show the dynamics of sensitive and resistant cancer populations. The blue crosses indicate the measurement points along the treatment trajectory. The dynamic test schedule decreases the length of on-medication periods and thus delays the development of resistance. **TTP** - time to progression. **Mean int** - mean length of the intervals between tests.

**Figure A3.**
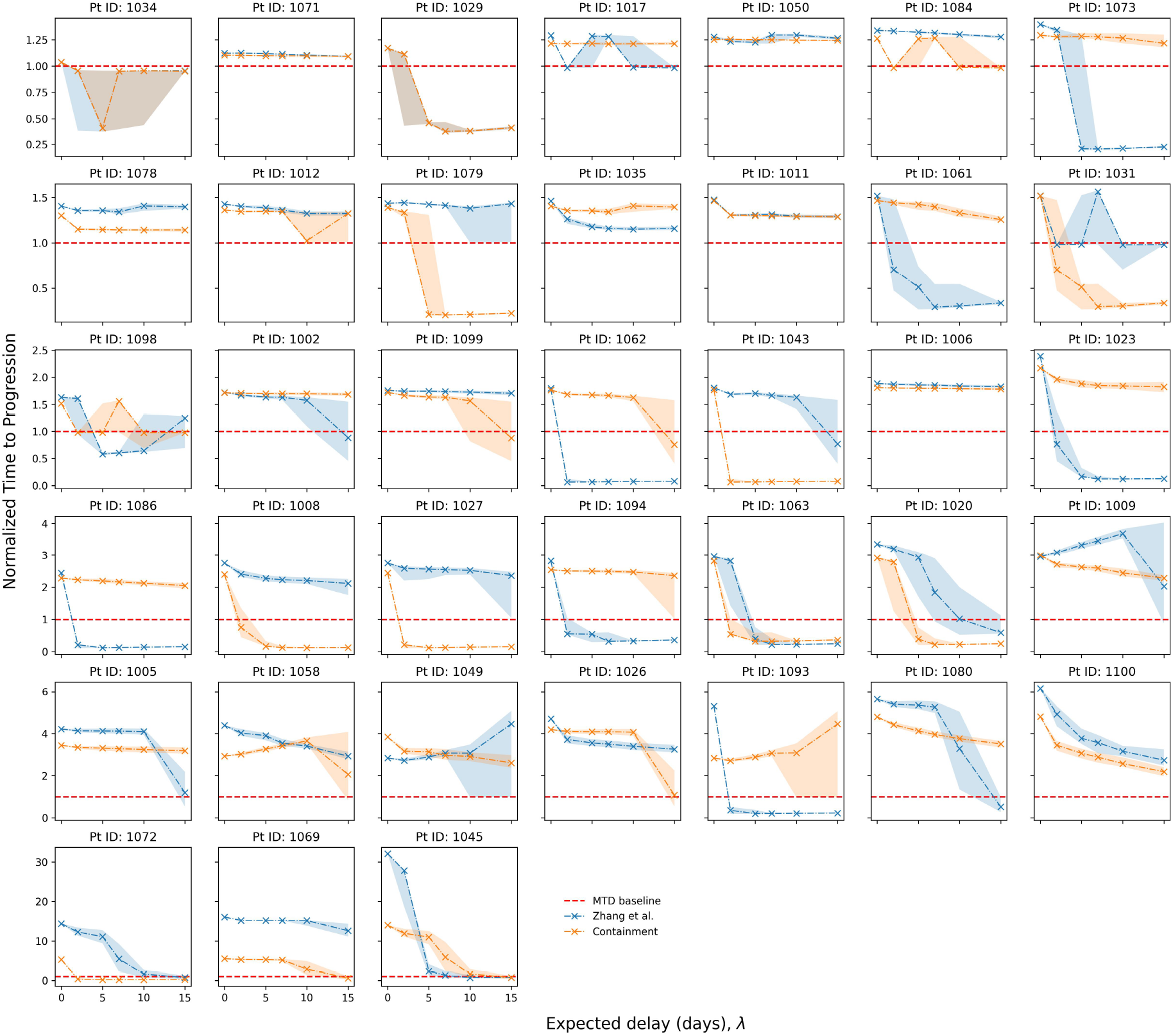
The dynamics of the median and interquartile range of TTPs with increasing expected appointment delay for 38 virtual patients under both Zhang et al.’s and containment protocol. The blue crosses indicate the median TTP of 1000 simulation runs under Zhang et al.’s protocol. The blue-shaded area shows the interquartile range. The orange crosses and shaded area show the same for the containment protocol. The TTPs are normalized by the MTD baseline - TTP under the MTD protocol without delay, which is indicated by the red dashed line.

**Figure A4.**
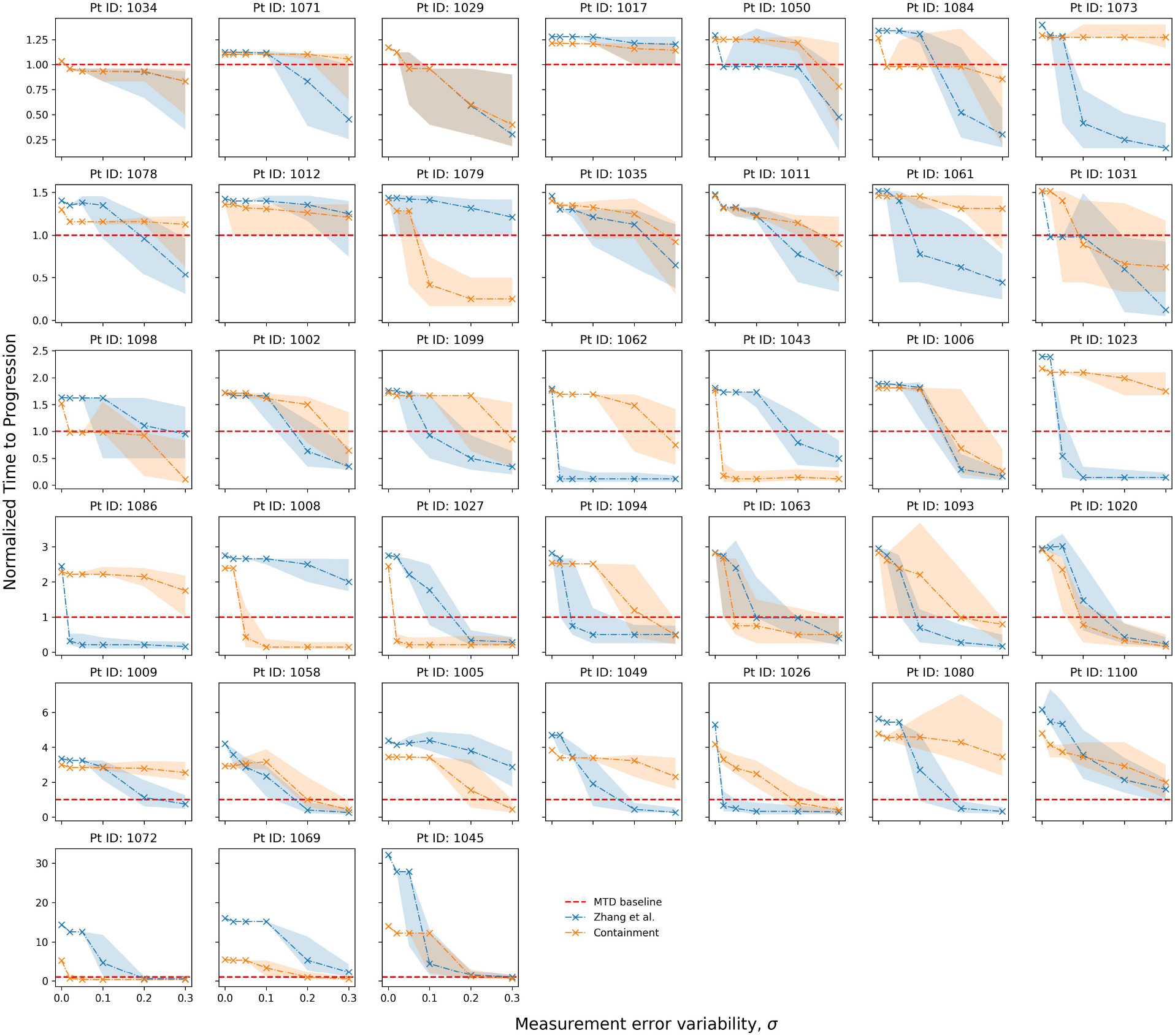
The dynamics of the median and interquartile range of TTPs with increasing measurement error variability (*σ*) for 38 virtual patients under both Zhang et al.’s and containment protocols. The blue crosses indicate the median TTP of 1000 simulation runs under Zhang et al.’s protocol. The blue-shaded area shows the interquartile range. The orange crosses and shaded area show the same for the containment protocol. The TTPs are normalized by the MTD baseline - TTP under the MTD protocol without error, which is indicated by the red dashed line.

**Figure A5.**
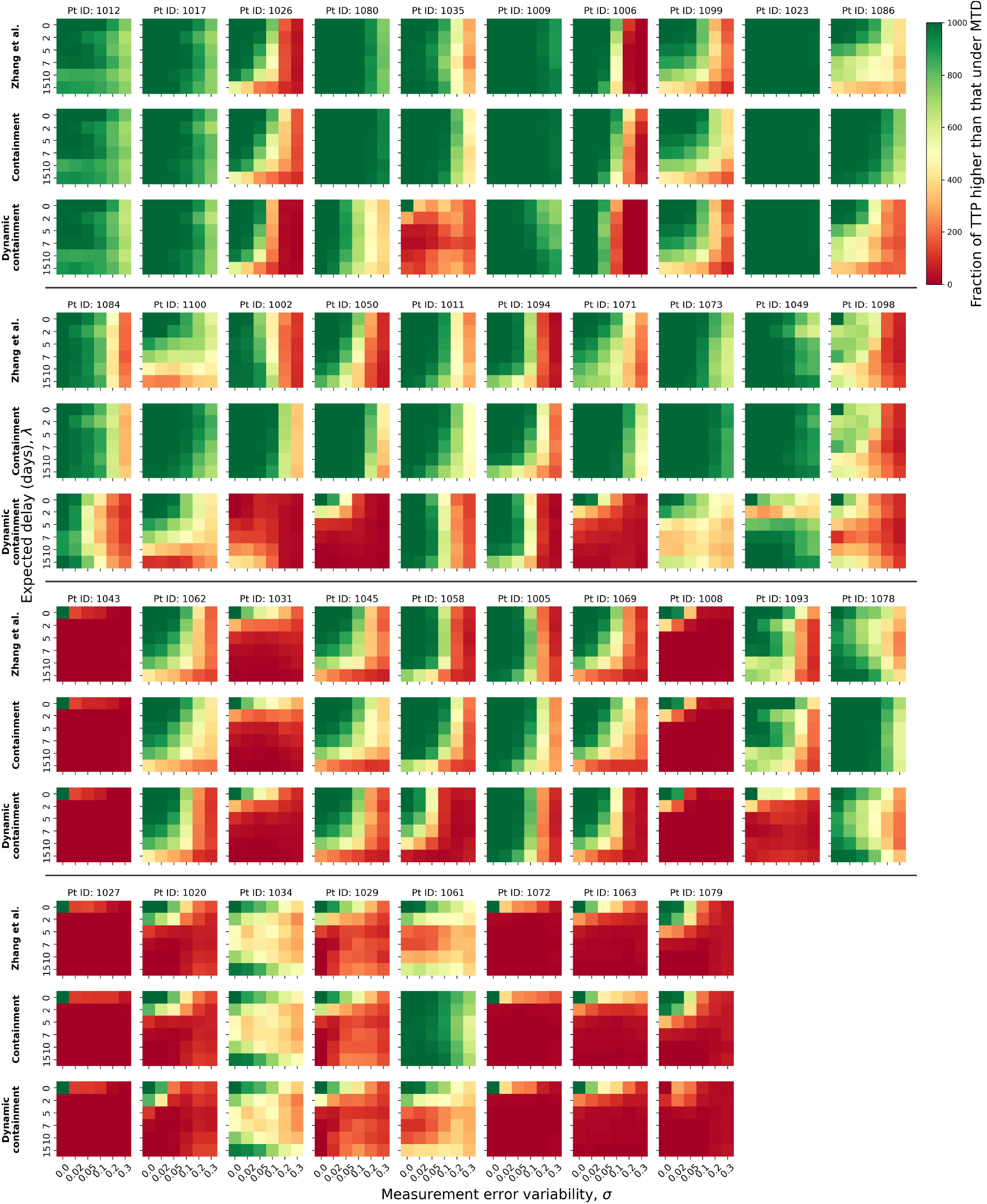
Heatmaps of the fraction of the simulation runs resulting in TTP higher than the MTD baseline under Zhang et al.’s protocol, containment and dynamic containment protocols for 38 virtual patients. The virtual patients are sorted in accordance with increasing growth rate in the absence of medication (Figure A7).

**Figure A6.**
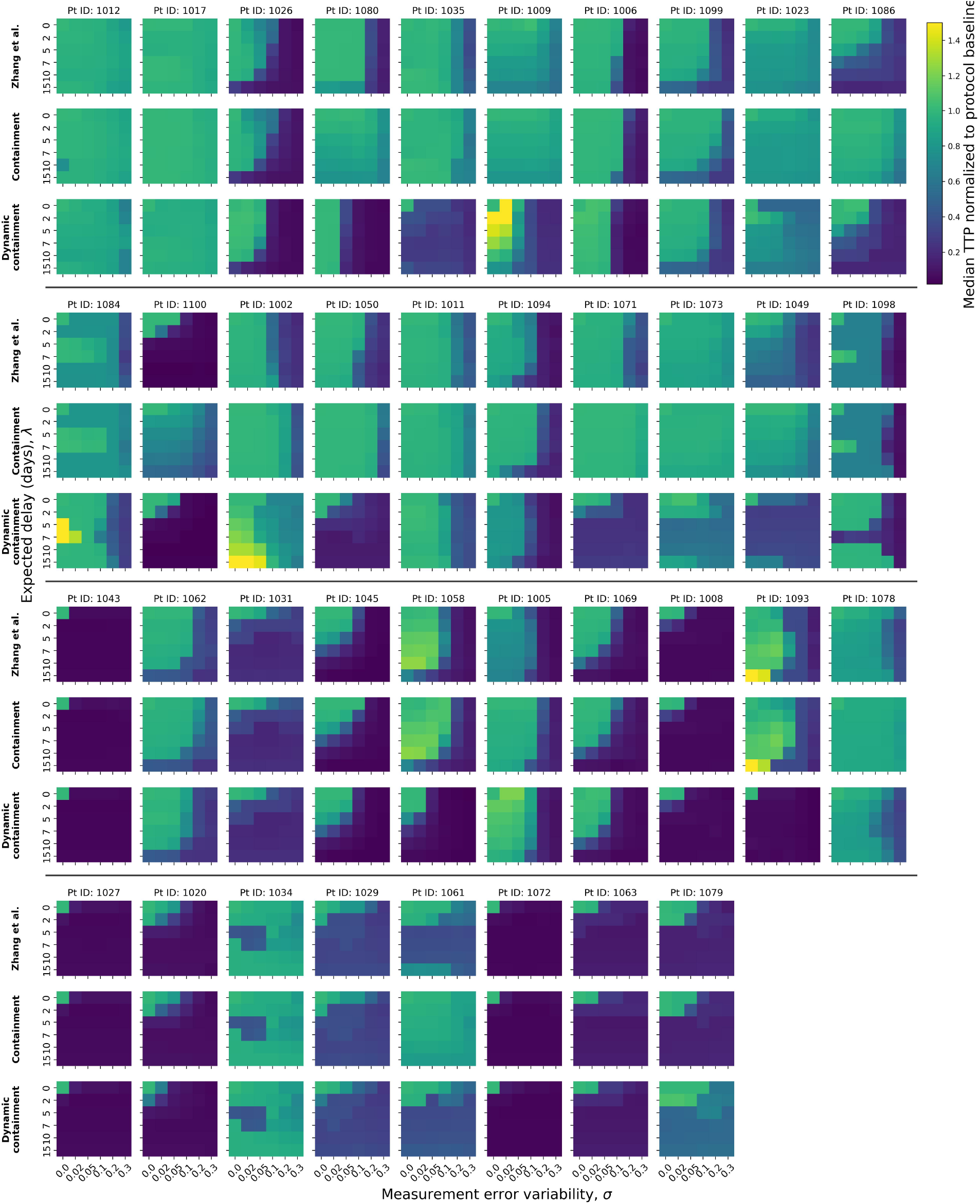
The change in median TTP with increase of measurement error variability (*σ*) and expected appointment delay (*λ*) normalized to the protocol baseline (TTP in no error, no delay conditions) under Zhang et al.’s, containment and dynamic containment protocols for 38 virtual patients. The virtual patients are sorted in accordance with increasing growth rate in the absence of medication (Figure A7).

**Figure A7.**
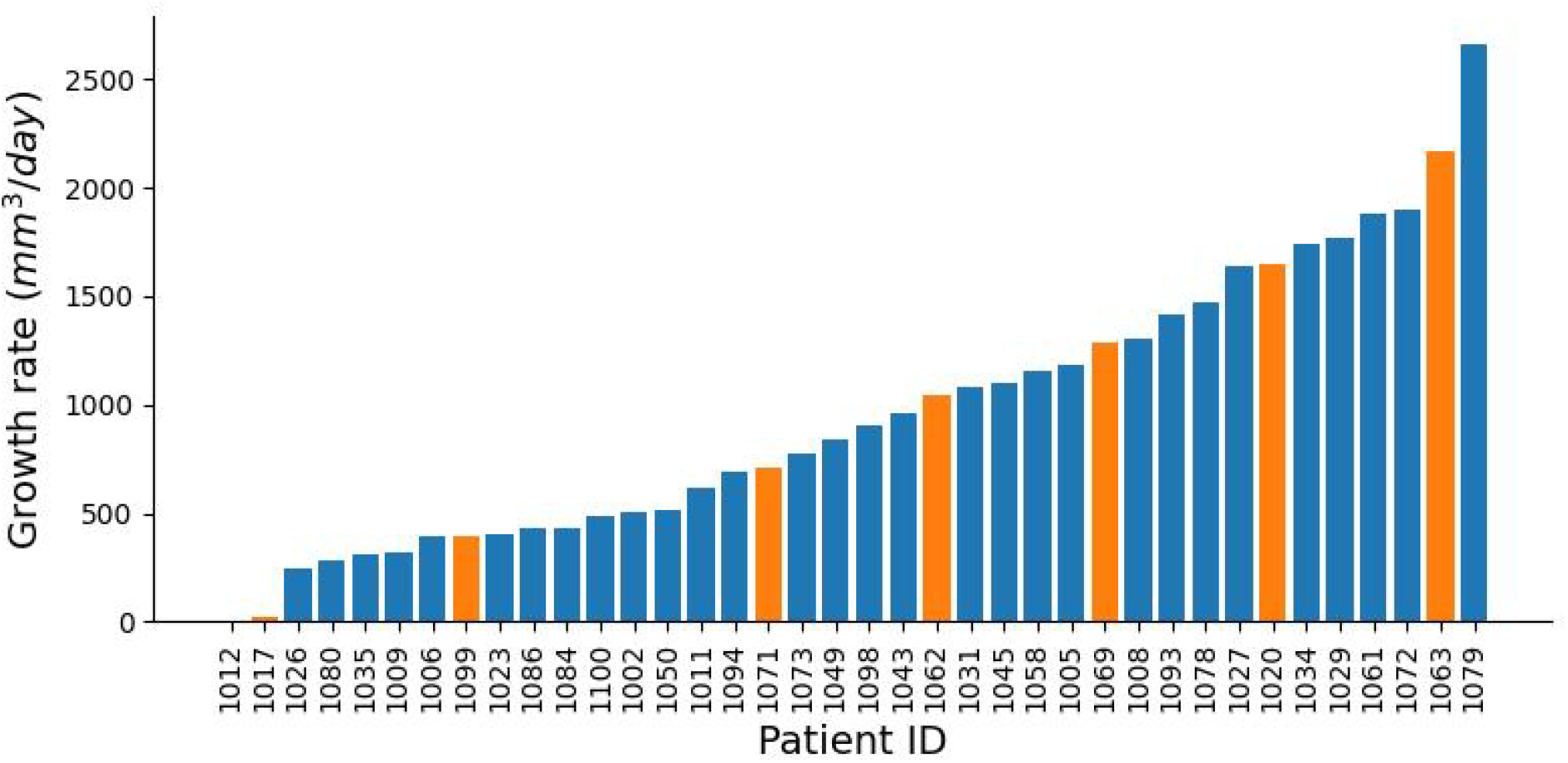
The linear growth rate of 38 virtual patients without treatment. The linear growth rate was estimated by simulating tumor volume growth in the absence of treatment at the start of therapy. The patients marked in orange are those presented in Figure 7.

**Table A1.** Percentage of virtual patients for whom the median TTP is higher under Zhang et al.’s protocol with the constant test schedule than with the dynamic schedule (left) and for whom the median TTP is higher under the containment protocol than under Zhang et al.’s protocol with the dynamic schedule (right) across delay and error combinations. The left part of the table shows the percentage of virtual patients for whom the median TTP under Zhang et al.’s protocol with a constant schedule was higher than that with a dynamic schedule. The right part of the table shows the percentage of virtual patients for whom the median TTP under the containment protocol was higher than under Zhang et al.’s protocol with a dynamic schedule. The rows indicate the mean delay *λ* in days, and the columns - the measurement error variability *σ*. Zhang et al.’s protocol with a constant schedule yields the highest TTP for the majority of patients with little or no error, but if the error is large, the protocol with a dynamically adjusted schedule outperforms the constant schedule, though not the containment protocol.

| Zhang et al. vs dyn sched Zhang et al. |  |  |  |  |  |  | Containment vs dyn sched Zhang et al. |  |  |  |  |  |  |
| --- | --- | --- | --- | --- | --- | --- | --- | --- | --- | --- | --- | --- | --- |
| $\lambda$ | $\sigma$ | | | | | | $\lambda$ | $\sigma$ | | | | | |
|  | 0.00 | 0.02 | 0.05 | 0.10 | 0.20 | 0.30 |  | 0.00 | 0.02 | 0.05 | 0.10 | 0.20 | 0.30 |
| **0** | 74 | 66 | 50 | 47 | 39 | 32 | **0** | 45 | 34 | 24 | 26 | 79 | 82 |
| **2** | 76 | 71 | 68 | 39 | 26 | 32 | **2** | 34 | 26 | 29 | 29 | 74 | 82 |
| **5** | 74 | 76 | 68 | 39 | 39 | 32 | **5** | 39 | 37 | 42 | 42 | 79 | 74 |
| **7** | 68 | 68 | 58 | 39 | 39 | 32 | **7** | 42 | 42 | 42 | 45 | 74 | 82 |
| **10** | 71 | 66 | 63 | 55 | 37 | 34 | **10** | 45 | 42 | 42 | 50 | 71 | 74 |
| **15** | 58 | 63 | 53 | 50 | 55 | 37 | **15** | 47 | 50 | 45 | 47 | 74 | 76 |

